# A Genome-wide CRISPR screen unveils the endosomal maturation protein WDR91 as a promoter of productive ASO activity in melanoma

**DOI:** 10.1101/2024.03.29.587208

**Authors:** Grégory Menchon, Aris Gaci, Antti Matvere, Marc Aubry, Aurélien Bore, David Gilot, Aurélie Goyenvalle, Rémy Pedeux

## Abstract

Antisense oligonucleotides (ASOs) belong to promising therapeutics for the treatment of neurologic, muscular and metabolic disorders. Several ASOs have been approved so far and more than a hundred clinical trials are currently underway covering a dozen therapeutic areas. Yet, the mechanisms of internalization and cell trafficking of these molecules remain poorly understood. Moreover, with only a small fraction of ASOs reaching the correct cellular compartment following systemic delivery, the majority of targeted diseases requires recurrent injections of ASOs. A deeper understanding of these mechanisms would guide the improvement of their potency and thus, reduce the amount of delivered ASOs and their potential side-effects. Here, using a CRISPR screen, we investigated intracellular proteins involved in ASOs efficiency using a whole genome approach and identified several potential regulators which could significantly impact ASOs potency in melanoma cells. We validated WD Repeat Domain 91 (WDR91), a regulator of endosomal maturation, as a modulator whose depletion significantly inhibits ASO productive activity. This study provides the first list of ASO modulators using a biologically relevant assay to estimate the role of these proteins. In conclusion, these data could lead to a better understanding of the mechanisms favoring productive uptake or improved endosomal escape of ASOs.

## INTRODUCTION

Antisense Synthetic Oligonucleotides (ASOs) belong to a class of attractive therapeutic compounds for numerous diseases. ASOs modulate cellular target gene expression through Watson-Crick binding to complementary RNA molecules and, according to their type, act by either promoting a RNAse-H1 mediated RNA degradation (Gapmers - GMs) or through a steric blocking mechanism (mixmers and target site blockers – TSBs) leading to three major modes of action: inhibition or activation of gene-expression and splicing modulation.^1^ ASOs have been developed for the treatment of different genetic disorders and they can be translated to personalized therapies.^2^ ASOs are specially taken up by fast growing cells^3^ and, among numerous advantages, bypass the current bottleneck regarding the targets that are considered undruggable with traditional small molecule inhibitors.^4^ Several ASOs are currently approved by the Food and Drug Administration (FDA) for genetic and rare diseases^5,6^ and several phase II and III studies are ongoing in the oncology therapeutic area.^7^

Although ASOs promise a great leap forward in future cancer treatments, their utilization suffers from major hurdles. Among them, a lack of tissue specific targeting,^8,9^ poor cellular uptake^10^ and the need to escape from the endo-lysosomal pathway to exert their role in either the cytoplasm or nucleus of the cells.^11,12^ Moreover, the mechanisms of internalization, cell trafficking and action of these molecules remain poorly understood. A small fraction of ASO reaching the right cellular compartment and target following systemic delivery^13,14^ leads to treatments consisting of high doses, recurrent injections and thus, significant accumulation in other non-targeted cells and organs with consecutive toxicities.^15^ It is therefore essential to better understand the fate of internalized ASOs and their intracellular route and mode of action. Such deeper characterization in different cell lines and with different ASO types and chemistries would help in the rational design and improvement of therapeutic ASO strategies in future clinical applications. Several studies have demonstrated that the fate of internalized phosphorothioate (PS)-containing ASOs was determined by intracellular interacting proteins.^16,17,18^ Even if a clear mechanism is still to be fully understood, around 80 intracellular proteins have been identified and this protein interactome appears to influence the pharmacological activities and potential toxicities of PS-ASOs by acting on their uptake and distribution. It is also worth noting that many other proteins that bind to ASOs have no effect on their activity. PS-ASOs are internalized and processed through the endocytic pathway,^3,19^ but proteins can modulate endosome traffic and maturation without interacting directly with oligonucleotides. Previous studies have investigated the modulation of intracellular pathways to increase the productive delivery and activity of oligonucleotides and to further unveil the critical molecular factors that contribute to nucleic acids pharmacological effects.^20^ Among these, modulation of endocytic recycling,^21,22^ multivesicular bodies’ (MVBs) fusion with lysosomes,^23^ endosomal escape^24^ and nuclear shuttling^25,26^ were explored. All these strategies involve the use of either genetic or non-genetic perturbations and have proven to be successful in many *in vitro* and *in vivo* studies. ASO-interacting proteins have been mostly characterized by *in vitro* studies using pull-down experiments on cell lysates^27,28^ but such method lacks physiological relevance as the intracellular membrane’s integrity is not preserved and could lead to many binding artefacts. Furthermore, pull-down studies may highlight some important ASOs protein modulators but only allow the identification of direct interactants. To circumvent these issues other authors recently used proximity biotinylation assay to highlight oligonucleotides interactome in living cells and in pharmacologically relevant conditions.^29^

Clustered Regularly Interspaced Short Palindromic Repeats (CRISPR) screen is a powerful genetic perturbation method that can be used for functional genomic studies, by activating or silencing genes.^30^ Then, it can be implemented to identify ASOs direct and indirect modulators in a more physiological state and preserving the cell membrane’s integrity. This technique was implemented in different studies to investigate modulators of dsRNAs,^31^ Antibody-Drug Conjugates (ADCs)^32^ or encapsulated mRNAs,^33^ but to our knowledge, only one study describes such a functional genomic screen (CRISPR gene activation) to uncover factors enhancing ASOs activity.^34^

In the present work, we used a genome-wide CRISPR screen strategy (CRISPR knockout), to uncover proteins that modulate the potency of an anti-proliferative tricyclo-DNA (tcDNA) based-ASO in a melanoma cell line (501Mel). The generation of the parental knockout cell library was performed such as every single cell had a unique depleted coding gene. The library was then treated with ASO, and sequencing was performed on the surviving (enriched or depleted) cells. This analysis highlighted several candidates of interest (95 activators and 55 inhibitors). From the top candidates, the robustness of our assay also allowed to highlight the proteins Ras-related protein Rab5C (Rab5C) and Component of Oligomeric Golgi Complex 8 (COG8) which are known positive regulators of ASOs productive trafficking and activity.^35,36^ We unveiled the endosomal maturation protein WDR91 as a strong positive regulator of the tcDNA based-ASO activity in this cell line. Activity modulation by WDR91 was validated together with Annexin A2 (ANXA2), a well characterized endosomal protein which promotes ASOs trafficking and productive activity in different cell lines.^37^ The impact of WDR91 was validated using two different ASO chemistries but was not validated on a splice switching ASO in a muscle cell line, highlighting a cell type-dependent effect. Finally, this work provides a detailed source of potential positive and negative regulators of ASO activity, coming from an unbiased screen and with a strong physiological relevance.

## RESULTS

### Development and *in vitro* validation of a tcDNA-ASO to inhibit 501Mel cells proliferation with a target-site blocking mechanism

Previous work aiming at masking a microRNA (miRNA) target site on a microRNA sponge (*TYRP1* mRNA) led to the test and validation of a commercially-designed Locked Nucleic Acid (LNA) TSB (named TSB-T3 for Target-Site Blocking – T3). In this study, TSB-T3 significantly induced the reduction of melanoma tumor growth *in vitro* and in 501Mel xenografts.^38^ This ASO was validated in five different cell lines and had a dose-response anti-proliferative activity. TSB-T3 exerts its ASO masking activity by preventing the sequestration of the miR- 16 tumor suppressor by *TYRP1* mRNA and redirecting it towards its target-RNAs, to induce their decay and subsequently reduce tumor proliferation (**Figure 1A**).^39^ This miRNA displacement mediated by a Target Site Blocker ASO represents a relevant assay to investigate the proteins involved in ASOs efficiency. The ASO blocking activity is directly monitored through an anti-proliferative effect, which has been validated *in vivo*.

**Figure 1.**
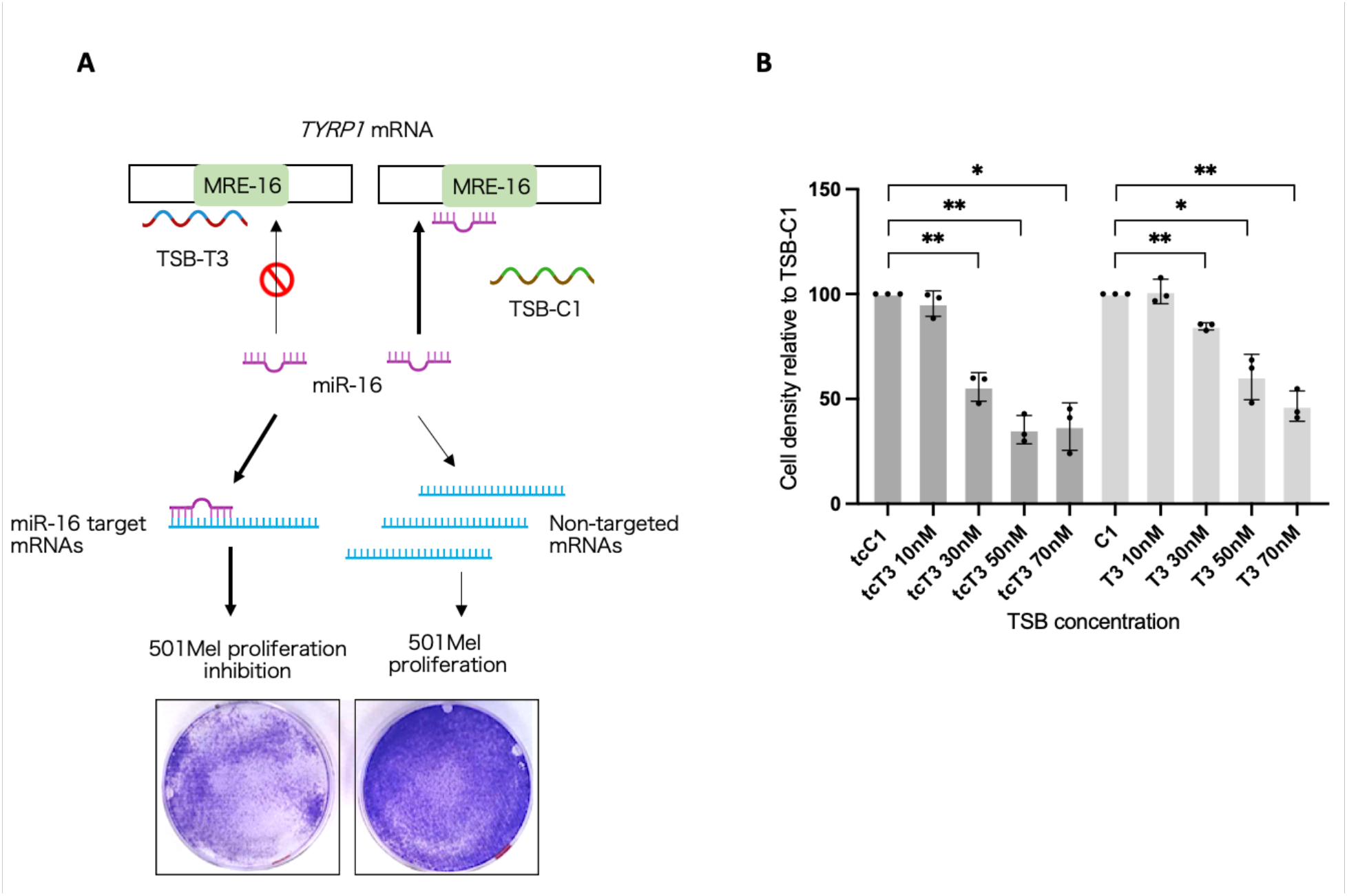
Inhibition of 501Mel cells proliferation with Target Site Blocker ASOs targeting *TYRP1* mRNA. (A) A competition mechanism occurs with binding of TSB-T3 to a sequence overlapping the miR-16 site in the 3’UTR of *TYRP1* mRNA, thus releasing miR-16 whose tumor suppressor activity is restored. TSB-C1 (Control ASO) has no match with *TYRP1* 3’UTR.^38^ The cell density assay upon TSB-C1 or -T3 treatment is measured through a crystal violet colorimetric assay. The figure 1A was generated with BioRender.com (B) 501Mel cell density assay at day 3 with increasing concentrations of reverse-transfected LNA- (T3) or tcDNA- (tcT3). Experiments were performed in independent biological triplicates. Data were normalized to the corresponding control conditions (C1 or tcC1) at the same concentrations (10, 30, 50 and 70nM). Data are presented as mean (SD). Unpaired *t*-test with Welch’s correction were performed. The statistical significance is **P=0,0040 (T3 *vs* C1 – 30nM); *P=0,0241 (T3 *vs* C1 – 50nM); **P=0,0061 (T3 *vs* C1 – 70nM); **P=0,0078 (tcT3 *vs* tcC1 – 30nM); **P=0,0036 (tcT3 *vs* tcC1 – 50nM); *P=0,0105 (tcT3 *vs* tcC1 – 70nM).

In the present study, we decided to look for protein modulators of an ASO with a clinically relevant chemistry. We thus designed and evaluated a new TSB-T3 with the same sequence but with a full tcDNA chemistry and phosphorothioate (PS) backbone (tcT3). This class of conformationally constrained ASO displays enhanced binding properties to DNA and RNA as well as unique pharmacological properties and unprecedented uptake by many tissues after systemic administration.^40,41^

To compare both chemistries, 501Mel cells were reverse-transfected in 96-well plates with either the LNA- or the tcDNA-ASO (named respectively T3 and tcT3). We deliberately used high doses of ASOs (up to 70nM) as their blocking activity is based on the displacement of a sequestered miR-16 by *TYRP1* mRNA which is, besides, highly expressed in melanoma cells (with around 3200 copies per cell).^38^ After 72h, the cell density was measured by colorimetric assay and normalized to a control ASO treatment (C1 or tcC1). As expected, we confirmed previous results with the commercial reagent. Both T3 and tcT3 ASOs showed a dose-response anti-proliferative activity and, interestingly, the tcDNA chemistry improved the ASO potency at low doses (**Figure 1B**). This tcDNA-ASO was selected for the subsequent screening experiment.

Genome-wide pooled sgRNA library generation, knockout library treatment with tcDNA- ASOs and NGS sequencing The CRISPR-knockout cell library was generated by transducing 501Mel cells with a pooled lentiviral library which contains the Cas9 and sgRNAs targeting 19.114 human coding genes. This lentivirus library contains 4 different guides per coding gene (total of 76.441 sgRNAs) and 1000 non-targeting control sgRNAs. A multiplicity of infection (MOI) of 0.4 was selected to ensure that only few cells are transduced with more than one sgRNA and avoid unspecific screening results.^42^ With 76.441 sgRNAs and a desired coverage of at least 400 cells per guide, a total of 32 million cells have been transduced and maintained during the screen.

After transduction, the cells were positively selected with antibiotic, expanded and subsequently transfected with either the tcT3 or -C1 ASOs. A final ASO concentration of 50nM was selected based on the former dose-response experiment which showed that the anti-proliferative effect reached a plateau at this dose (**Figure 1B**). The treatment was performed for three days. After this period, the difference in cell density between the treated and control conditions was greater than 50%.

At the end of treatment, the cells were expanded, pooled and a pellet of 100 million cells per condition was used for genomic DNA (gDNA) extraction and PCR amplification of the integrated sgRNAs in the 3 conditions (non-treated parental cell bank, tcT3 treated and tcC1 treated). The global workflow is depicted in **Figure 2A**.

**Figure 2.**
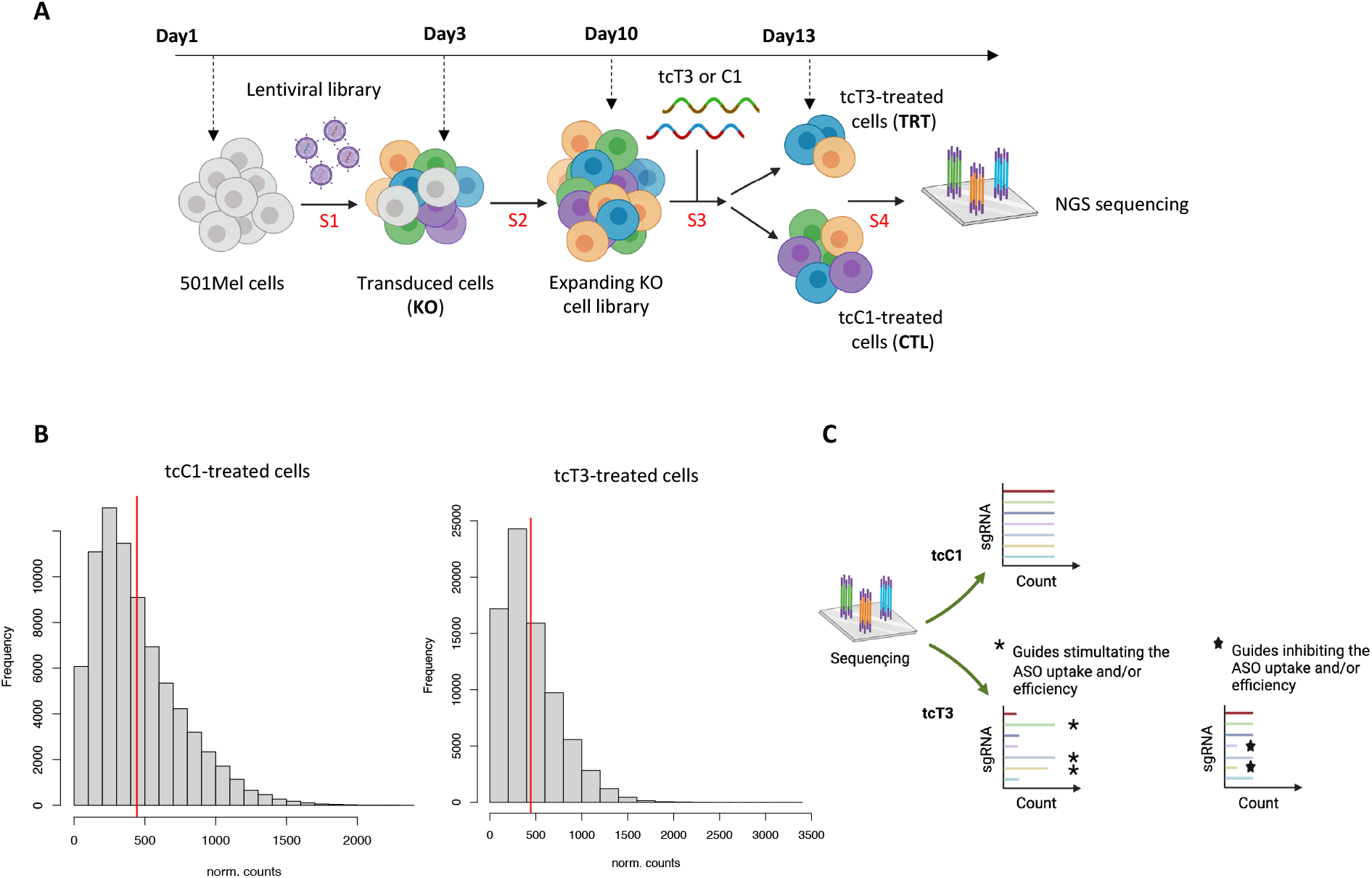
Generation and treatment of the 501Mel cells knockout library. (A) Workflow of the experimental setup. S1: Step 1 (cells transduction – non infected cells in grey on the workflow); S2: Step 2 (transduced cells positive selection with antibiotic); S3: Step 3 (ASO exposure); S4: Step 4 (gDNA purification and sgRNAs amplification). The figure was generated with BioRender.com (B) Distribution of the number of normalized read counts per sgRNA in the tcT3 and the tcC1-treated library. The mean is represented by a red bar (C) Method for identifying genes promoting or inhibiting ASO uptake and/or efficiency after NGS sequencing and based on normalized read counts after tcC1 or tcT3 treatment.

After amplicons sequencing by NGS, a quality check was performed and only 115 sgRNAs were not detected over 76.441 in our cell library (0.15%), which validated proper conditions before ASOs treatment. The normalized reads count in each control and therapeutic ASO condition were plotted and provided a mean of 400-500 reads per guide as expected (**Figure 2B**).

The sequenced sgRNAs coming from the pool of surviving cells after ASO treatment provided information regarding the enriched or depleted ones. Thus, by comparing the sgRNAs enrichment or depletion in treated *versus* control condition and for each gene, we sought to identify proteins potentially involved in the regulation of ASO uptake, trafficking, release and guidance to their RNA-target in this melanoma cell line (**Figure 2C**).

### Identification of tcT3 ASO activity regulators in 501Mel cells

To identify proteins that could impact the tcT3 activity in 501Mel cell line, the NGS sequencing raw data accounting for the total number of sgRNA reads per gene and per sample were analyzed as follows. First, we used MAGeCK count to collect sgRNA read count information from the tcC1- and tcT3-treated cells sequencing files. Then, we used MA Plots to score and identify candidates based on the sgRNA read abundance (A value) and the read counts ratio between the treated and the control condition (M value: fold change - FC). M and A values were computed for each of the 4 sgRNAs targeting each individual gene (**Figure 3A**). From the MA plots, top activator and inhibitor candidates were extracted after using a defined M cutoff for each sgRNA per gene. The **figure 3B** represents MA plot examples from selected activator and inhibitor candidates with applied M cutoff and with 4 (GGTLC2) or 2 relevant sgRNAs (WDR91). Because of a short time of treatment (72h), a minimum FC of 1.5 was fixed between the tcT3 and tcC1 read counts to highlight the best hits (activator candidates: fold change tcT3 read counts *vs* tcC1 read counts ≥ 1.5; inhibitor candidates: fold change tcC1 read counts *vs* tcT3 read counts ≥ 1.5) (Candidates examples in **Figure 3C**). We opted for a short time of treatment to avoid guide selection bias.

**Figure 3.**
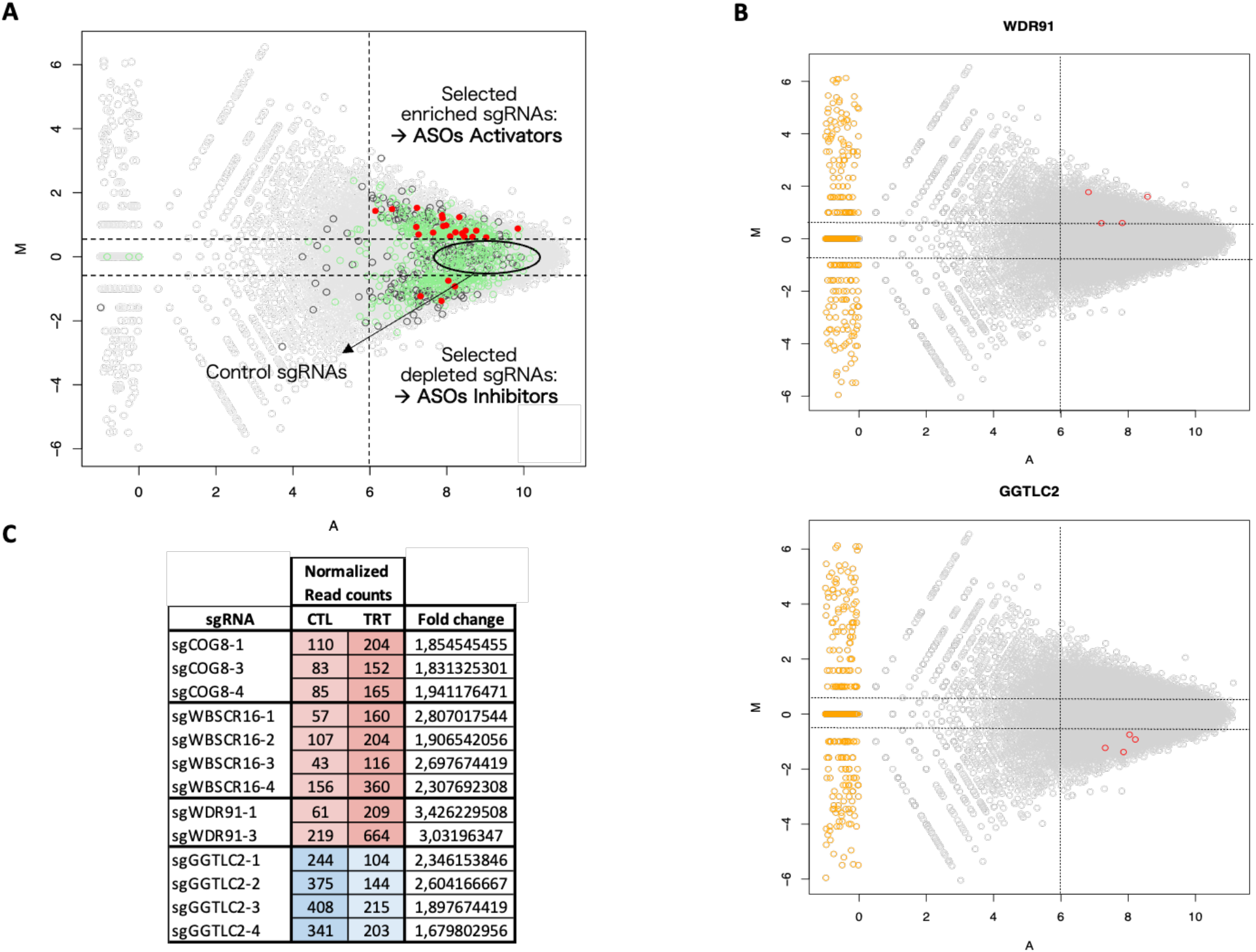
NGS sequencing data sorting and analysis. (A) After sgRNA extraction, MA plots are used to visualize the differences between measurements (read counts) between two conditions: treatment (tcT3) *versus* control (tcC1). For each sgRNA, the read counts are transformed onto M (log ratio) and A (mean average) scales. M is equal to log2(sgRNA_count_in_TRT+1) - log2(sgRNA_count_in_CTL+1) and A is the mean between count_TRT and count_CTL in log2 scale. Candidates’ selection was performed according to the following criteria: A ≥ 6 and M ≥ 0.6 (enriched sgRNAs, ASOs activators); A ≥ 6 and M ≤ -0.6 (depleted sgRNAs, ASOs inhibitors). Control sgRNAs (non-targeting guides) are located between M ≥ 0.6 and M ≤ -0.6. On the obtained MA plot, red, green and black dots represent the genes with the 4, 3 and 2 sgRNAs over 4 fulfilling the defined criteria, respectively (B) MA plot examples from selected activator and inhibitor candidates with applied M and A filters and with 4 (GGTLC2) or 2 relevant sgRNAs (WDR91). The yellow dots represent the genes with a very low expression in either the treatment or the control condition and as a consequence, an elevated but non-significant fold change (C) Representation of the post-sequencing normalized read counts in the control (CTL – tcC1) and treated (TRT – tcT3) conditions for the sgRNAs of four sorted modulator candidates (COG8, WBSCR16, WDR91, in red – activators; GGTLC2 in blue – inhibitor). The fold change is calculated as: tcT3 read counts / tcC1 read counts (for activators); tcC1 read counts / tcT3 read counts (for inhibitors).

Several sgRNAs targeting the same gene might not have a consistent behavior. However, the Brunello library has been designed with optimized sgRNAs and is known to outperform other knockout libraries.^43^ Indeed, the robustness of this library has already been demonstrated with the ability to identify reliable candidates with only 2 effective sgRNAs per gene.^43^ We thus extracted activator and inhibitor candidates considering two, three or four out of four guides meeting the defined M (FC) and A selection criteria.

As shown in **Figure 4A and B** (and in **Table 1**), this cutoff allowed to highlight a set of 95 activator candidates and 55 inhibitors. Regarding the activators, 5 genes had their 4 respective sgRNAs fulfilling the corresponding criteria; 28 genes had 3 effective sgRNAs and 62 genes had 2 effective sgRNAs over 4. From the inhibitors, 3, 7 and 45 genes had 4, 3 and 2 effective sgRNAs, respectively. Each of the candidates have been represented by an average FC value but a detailed table of each sgRNA and their respective M, A and FC values is available in **Table 1**.

**Figure 4.**
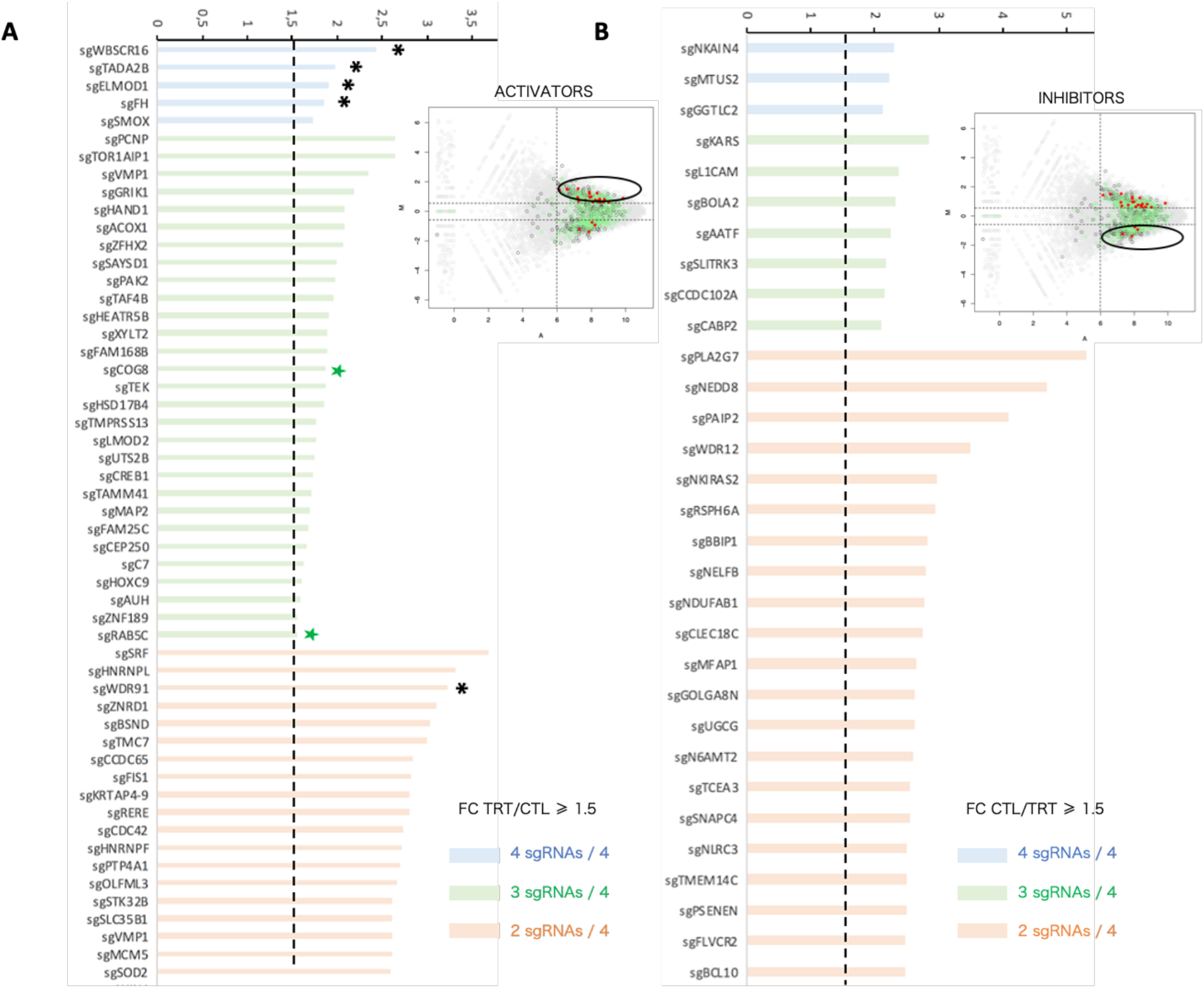
ASO activators and inhibitors ranking and selection. (A) Distribution of the average fold change (tcT3 read counts *vs* tcC1 read counts) from the fold changes of the 2, 3 or 4 corresponding sgRNAs per gene, for 53 of the 95 selected ASO activator candidates (fulfilling the criteria A ≥ 6 and M ≥ 0.6 on the MA plot). Green stars represent candidates that have already characterized in the literature. Black stars correspond to the 5 candidates that have been chosen for further functional validation (B) Distribution of the average fold change (tcC1 read counts *vs* tcT3 read counts) from the fold changes of the 2, 3 or 4 corresponding sgRNAs per gene, for 31 of the 55 selected ASO inhibitor candidates (fulfilling the criteria A ≥ 6 and M ≤ -0.6 on the MA plot). The fold change threshold was fixed to 1.5. The full list activator and inhibitor candidates is provided in **Table 1**.

**Table 1.**
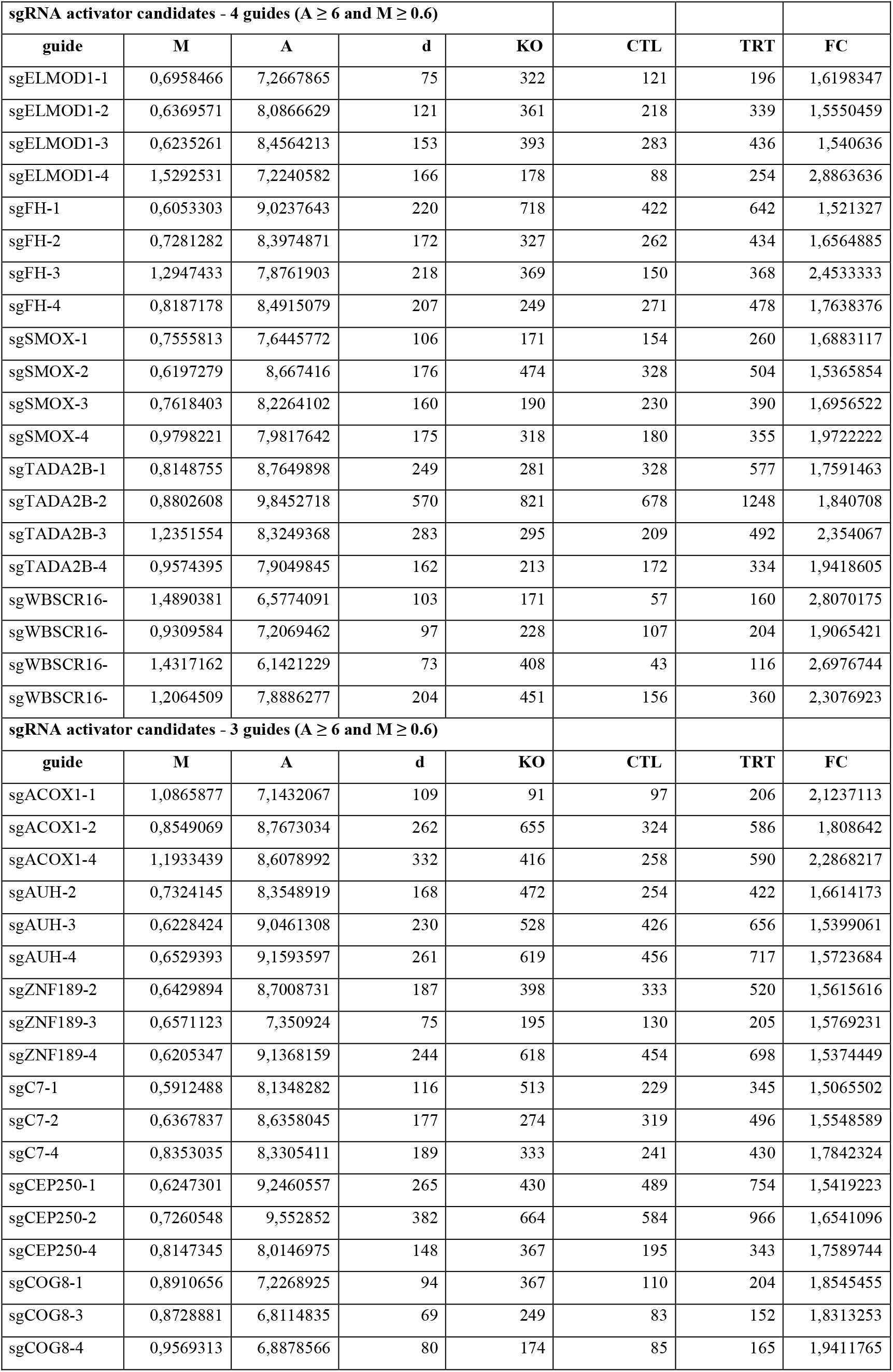
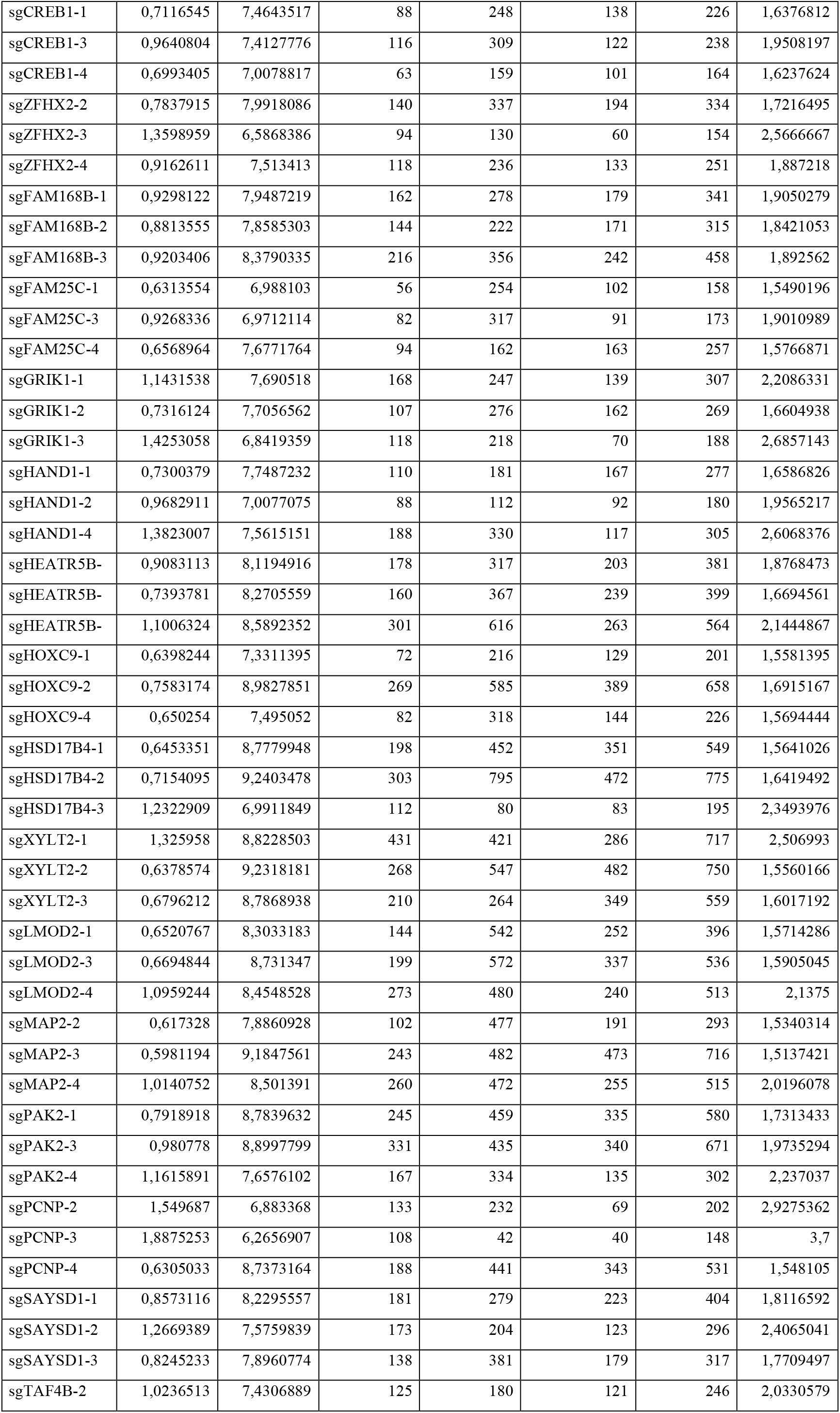
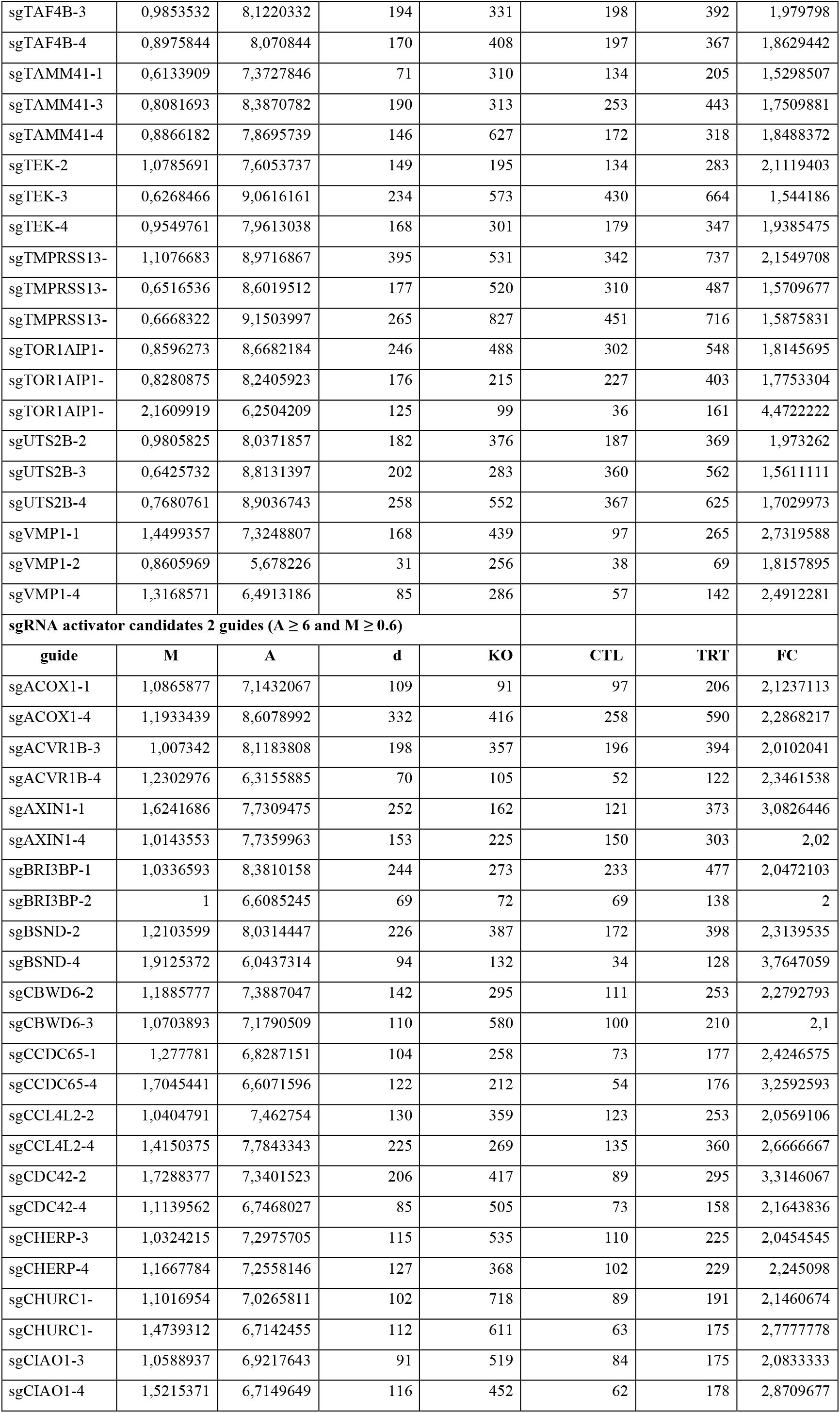
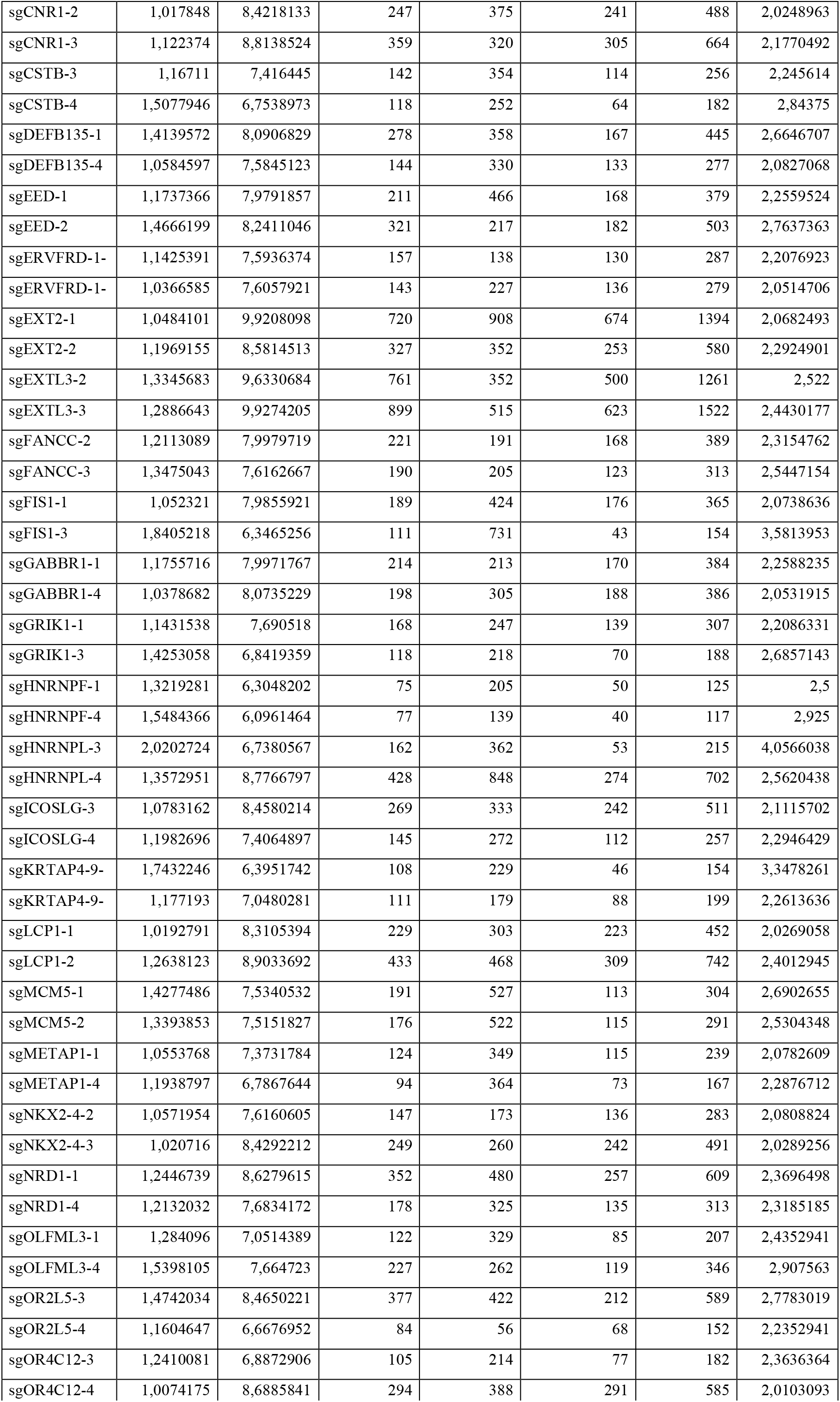
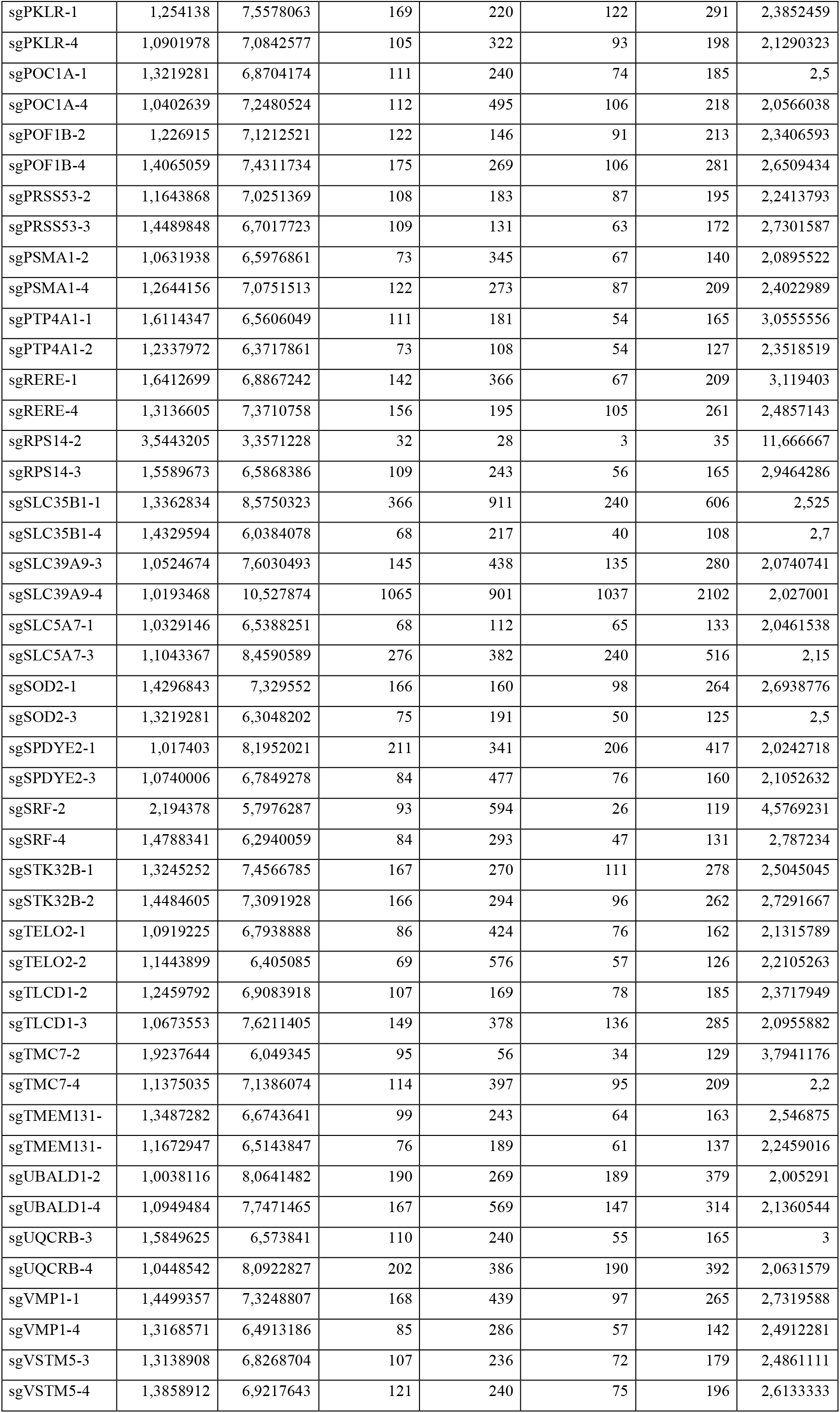
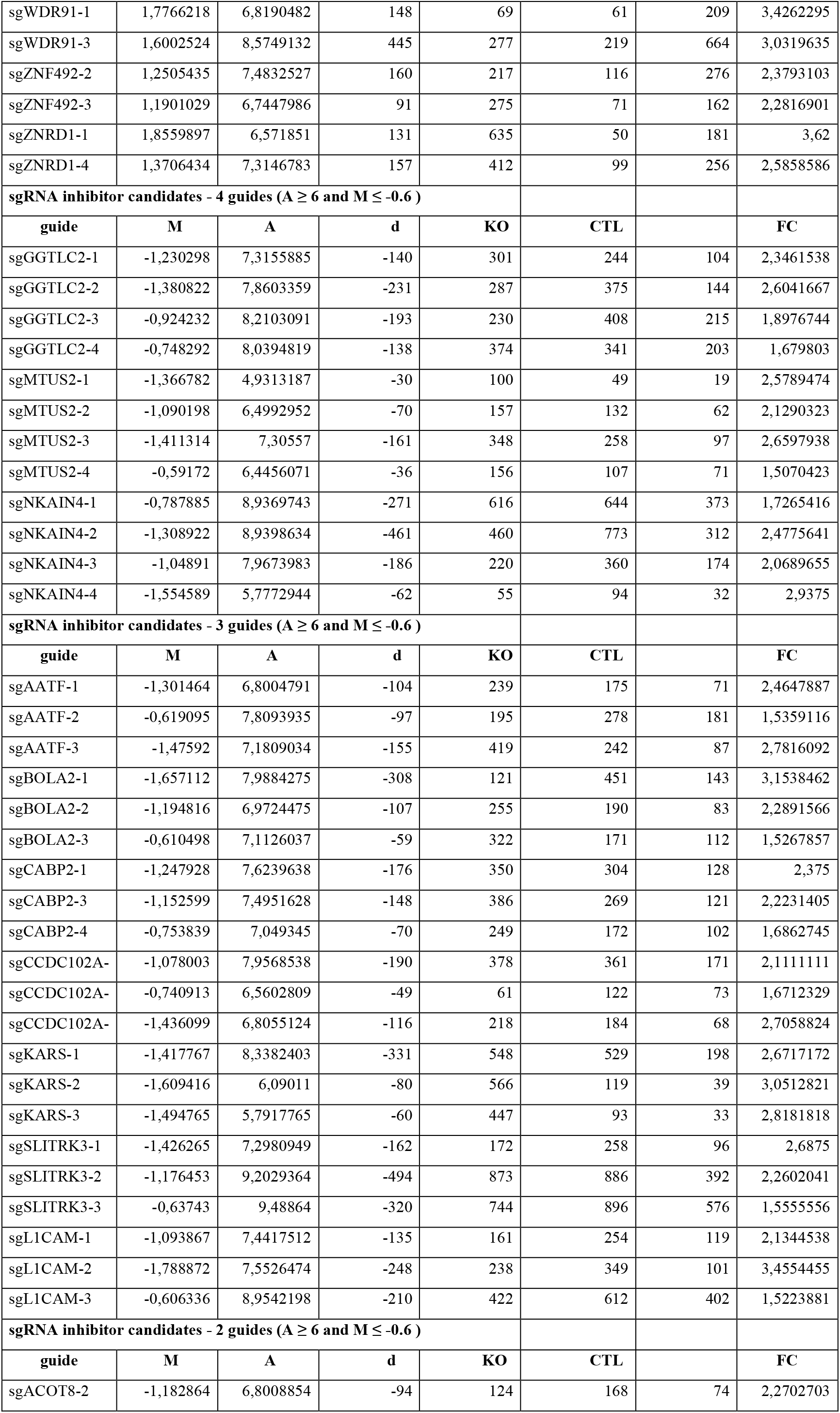
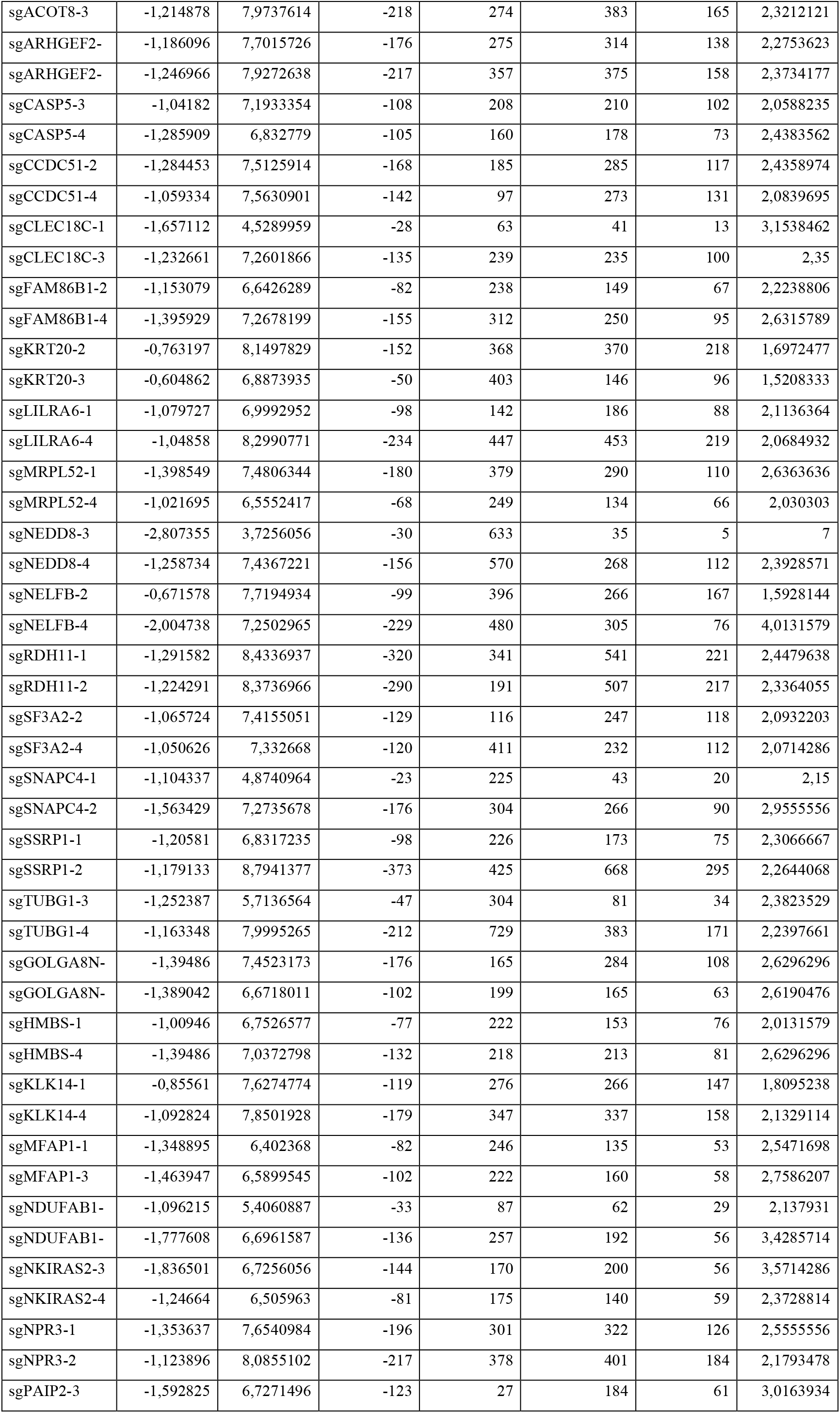
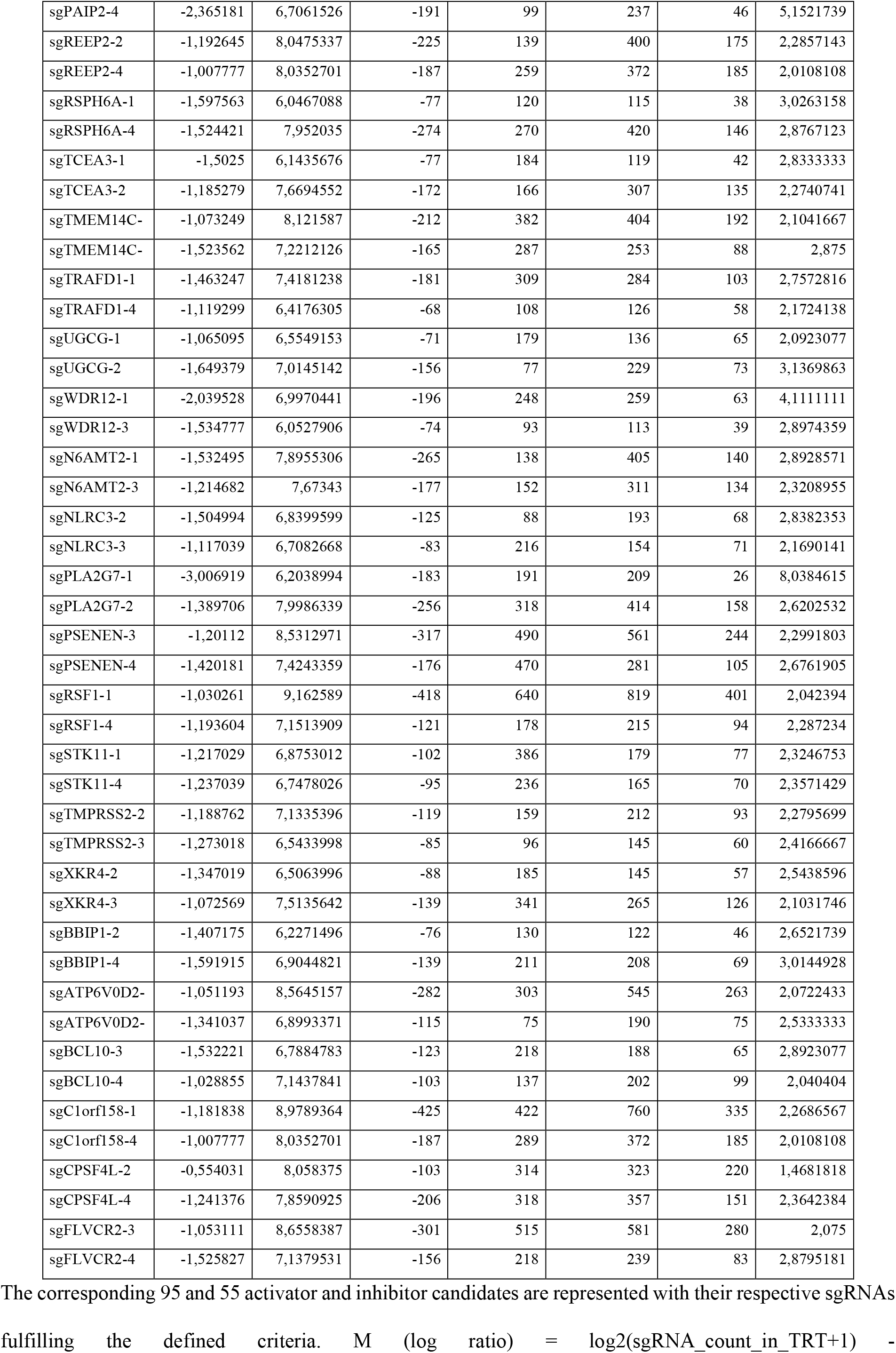
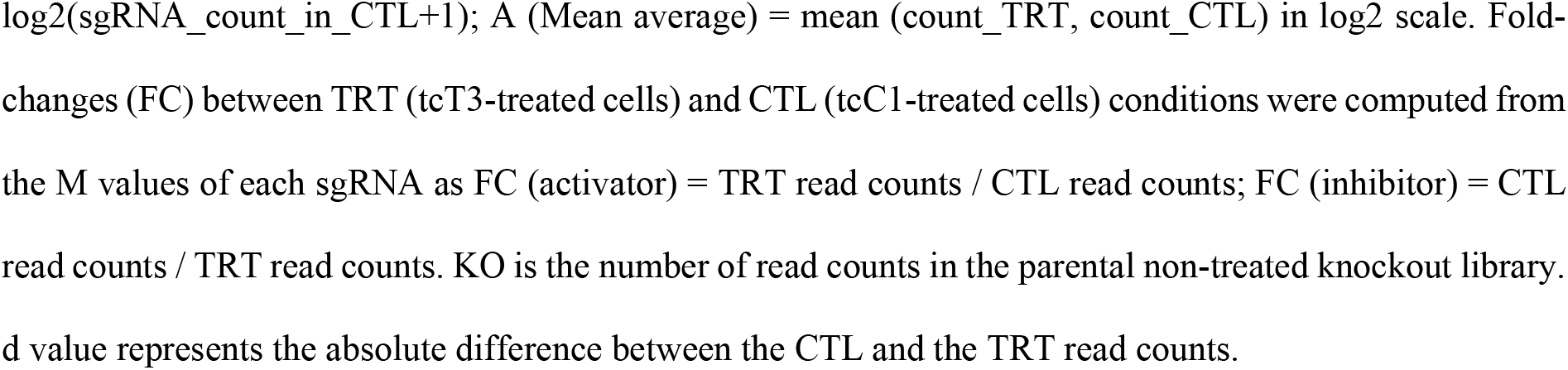
List of the ASO activator and inhibitor candidates sorted after applying M, A and fold change filters.

Gene set enrichment analysis using PANGEA (Pathway, Network and Gene-set Enrichment Analysis) allowed to classify the activator and inhibitor candidates based on the cellular component gene ontology (**Supplemental Figures S1 and S2**).^44^ Interestingly, the activator candidates with the highest fold enrichment are mainly clustered within proteins belonging to the peroxisome and mitochondria compartments. Inhibitor candidates are mainly clustered within proteins belonging to the Golgi apparatus and microtubules regulation.

For subsequent validation experiments, we decided to focus on protein activators of ASOs since we were more interested in the mechanistic uptake and trafficking of ASOs in cells. From the list of activators having 3 effective sgRNAs we highlighted COG8 and Rab5C, two known and well characterized positive regulators of PS-ASOs productive trafficking and activity in different cell lines.^35,36^ This finding further supports the validation and robustness of our screening experiment. As a first set of genes, we selected Elmo Domain Containing 1 (ELMOD1), Fumarate Hydratase (FH), Transcriptional Adapter 2-Beta (TADA2B) and Williams-Beuren Syndrome Chromosome Region 16 (WBSCR16), which are from to the top ranked candidates with 4 sgRNAs fulfilling the chosen criteria. We also selected WDR91 as it is an important factor that participates in the negative regulation of phosphatidylinositol–3– phosphate (PtdIns3P) allowing the early-to-late endosome conversion.^45,46^ This first pool of 5 genes was considered as a nonbiased sampling of potential activator candidates to be further investigated.

### WDR91 promotes tcT3- and LNA T3-ASOs activity in 501Mel cells

An RNA interference experiment was designed to further validate the five selected candidates as ASO activity enhancers (**Figure 5A**). After protein knockdown, the tcT3 activity should be reduced resulting in increased cell proliferation compared to a control condition.

**Figure 5.**
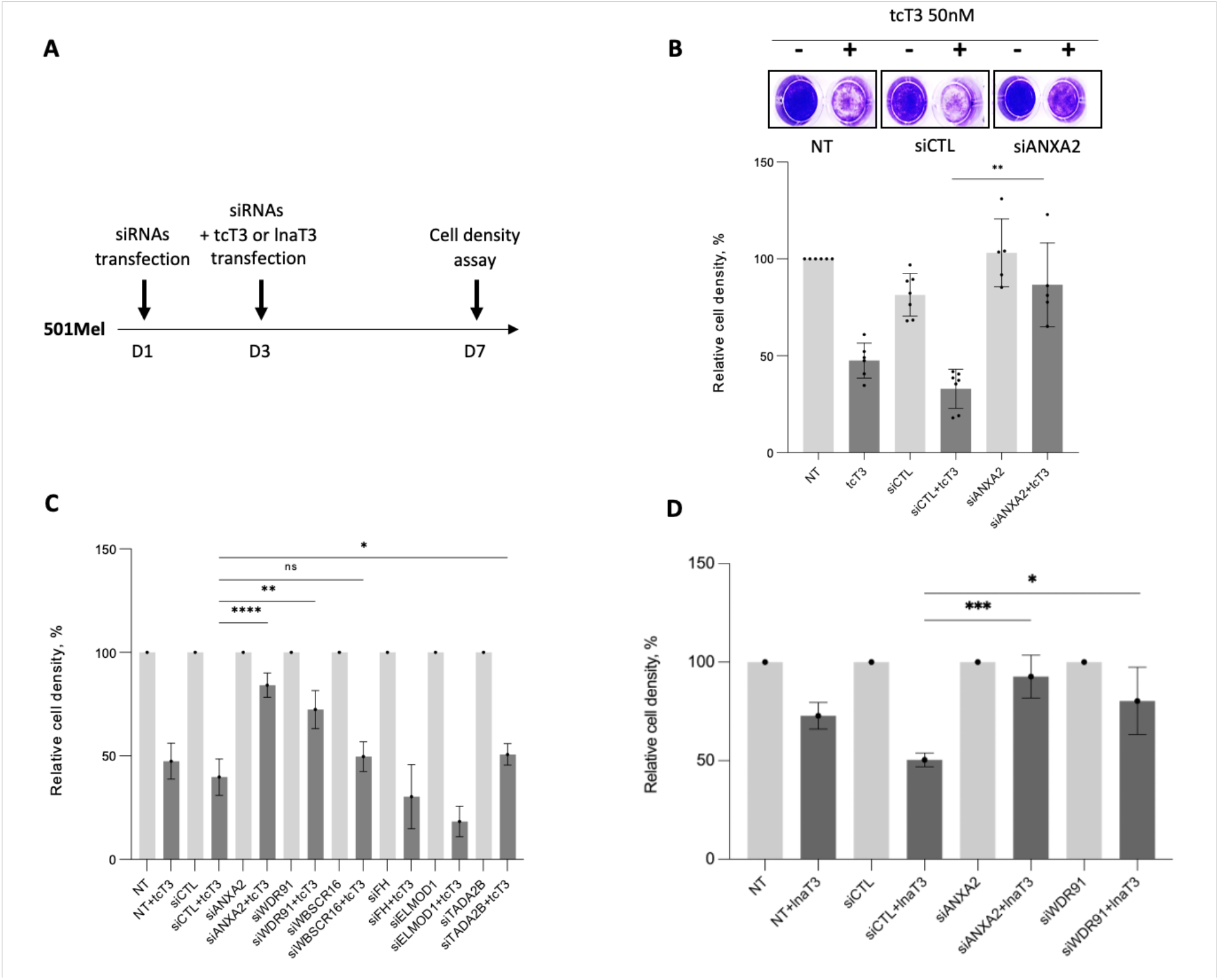
Functional validation of tcT3 activity enhancer candidates in 501Mel cells. (A) Workflow of the experimental design (B) Setup of the 501Mel cell density assay with ANXA2 positive control. Cell density measurement was performed by crystal violet assay after tcT3 transfection on normal, control or ANXA2 knockdown cells (NT: Non-Treated; siCTL: control siRNA). The relative cell density in each condition is normalized to the non-treated cells. (C) 501Mel cell density assay after tcT3 transfection on normal, control or candidate’s knockdown cells. Data are represented with a cell density normalized to 100% for each treatment with the siRNA alone, to discard any proliferation bias induced by the siRNA’s effects. (D) 501Mel cell density assay after LNA T3 transfection on normal, control, WDR91 or ANXA2 knockdown cells. Data are represented with a cell density normalized to 100% for each treatment with the siRNA alone, to discard any proliferation bias induced by the siRNA’s effects. Experiments were performed in independent biological replicates. Data are presented as mean (SD). Unpaired *t*-test with Welch’s correction were performed. The statistical significance is Panel C: *P=0,0468 (TADA2B, N=3); **P=0,0011 (WDR91, N=4); ****P≤0,0001 (ANXA2, N=5); NS= non-significant (P=0,0588); Panel D: *P=0,0154 (WDR91, N=5); ***P=0,0004 (ANXA2, N=5).

It was already well established in previous *in vitro* studies that the protein ANXA2 was an important facilitator of PS-ASOs endosomal trafficking in various cell lines.^37^ We thus first evaluated the effect of ANXA2 downregulation in 501Mel cells proliferation with or without tcT3 in comparison with a control siRNA. A western blot analysis confirmed the downregulation of ANXA2 protein after cells treatment with an ANXA2 siRNA (**Supplemental Figure S3A**). As shown in **Figure 5B**, *ANXA2* mRNA knockdown significantly impaired the activity of tcT3 by comparison with a control siRNA.

As for ANXA2, each individual candidate (WBSCR16, TADA2B, FH, ELMOD1 and WDR91) was downregulated with a corresponding siRNA (or a control siRNA) for 48h. The knockdown efficiencies were confirmed by qPCR analysis (**Supplemental Figure S3B** – Apart from ELMOD1, all knockdowns were shown to be significant enough to allow a relevant biological effect). 501Mel cells were then reverse transfected with either the siRNA alone or a combination of siRNA and tcT3. Four days after the second transfection, the cell density was measured by a crystal violet colorimetric assay (**Figure 5A**). After normalizing the cell density to each of the corresponding single siRNA treatment, WDR91 and TADA2B knockdowns showed a significant inhibition of tcT3 anti-proliferative activity (**Figure 5C and Table 2**). In these conditions, WDR91 knockdown cells showed the most significant and reproducible effect. Therefore, we decided to focus on WDR91 in our next investigations.

**Table 2.** Raw absorbance at 590nm (crystal violet coloration) of the cell density after four days of siRNA or siRNA + tcT3 ASO transfection.

|  | NT | NT+tcT3 | siCTL | siCTL+tcT3 | siWDR91 | siWDR91+tcT3 |
| --- | --- | --- | --- | --- | --- | --- |
| N1 | 2,251 | 0,781 | 1,534 | 0,403 | 2,644 | 1,585 |
| N2 | 2,93 | 1,53 | 2,62 | 1,19 | 3,01 | 2,39 |
| N3 | 2,89 | 1,42 | ND | ND | 2,98 | 2,42 |
| N4 | 2,6256 | 1,0660 | 1,8022 | 0,5007 | ND | ND |
| N5 | 2,8453 | 1,3494 | 2,518 | 1,19 | 2,927 | 2,022 |
| N6 | 2,7067 | 1,6489 | 2,229 | 0,9583 | ND | ND |
| N7 | 2,2459 | ND | 1,7195 | 0,8387 | ND | ND |
| N8 | 2,2341 | ND | 2,1565 | 0,8573 | ND | ND |

|  | siWBSCR16 | siWBSCR16+tcT3 | siFH | siFH+tcT3 | siELMOD1 | siELMOD1+tcT3 |
| --- | --- | --- | --- | --- | --- | --- |
| N1 | 2,829 | 1,4001 | 2,663 | 0,767 | 2,397 | 0,626 |
| N2 | 2,79 | 1,12 | ND | ND | ND | ND |
| N3 | 3 | 1,83 | ND | ND | ND | ND |
| N4 | ND | ND | ND | ND | ND | ND |
| N5 | ND | ND | ND | ND | ND | ND |
| N6 | ND | ND | ND | ND | ND | ND |
| N7 | 2,3493 | 1,2057 | 2,8127 | 1,3581 | 2,3740 | 0,4554 |
| N8 | 2,4994 | 1,162 | 2,3923 | 0,3284 | 1,8733 | 0,1822 |

|  | siTADA2B | siTADA2B+tcT3 | siANXA2 | siANXA2+tcT3 |
| --- | --- | --- | --- | --- |
| N1 | ND | ND | 2,95 | 2,77 |
| N2 | ND | ND | 2,69 | 1,91 |
| N3 | ND | ND | ND | ND |
| N4 | 1,3038 | 0,6641 | ND | ND |
| N5 | ND | ND | ND | ND |
| N6 | ND | ND | 2,3097 | 2,1038 |
| N7 | 1,4761 | 0,8328 | 2,3289 | 1,9428 |
| N8 | 1,663 | 0,7457 | 2,3167 | 1,8059 |

N1 to N8: biological replicates; NT: Non-Treated; siCTL: Control siRNA; ND: Not Determined.

We next wondered whether this mechanism was PS-ASO chemistry dependent and carried out the same experiment as above but with the LNA version of the TSB-T3. Again, in these conditions, ANXA2 and WDR91 knockdowns showed a significant inhibition of the LNA-T3 anti-proliferative activity (**Figure 5D and Table 3**), indicating that this mechanism was not PS-ASO chemistry dependent.

**Table 3.** Raw absorbance at 590nm (crystal violet coloration) of the cell density after four days of siRNA or siRNA + LNA T3 ASO transfection.

|  | NT | NT+lnaT3 | siCTL | siCTL+lnaT3 | siANXA2 | siANXA2+lnaT3 | siWDR91 | siWDR91+lnaT3 |
| --- | --- | --- | --- | --- | --- | --- | --- | --- |
| N1 | 2,91 | 1,9 | 1,78 | 0,82 | 2,42 | 1,75 | 2,69 | 1,65 |
| N2 | 2,95 | 2,04 | 1,91 | AM | 2,99 | 2,98 | 2,92 | 2,9 |
| N3 | 2,9 | 2,43 | 2,28 | 1,19 | 2,85 | 2,82 | 2,83 | 2,82 |
| N4 | 2,97 | 2,25 | 1,15 | 0,63 | 2,95 | 2,87 | 2,87 | 1,92 |
| N5 | 2,9 | 2,03 | 1,93 | 0,94 | 2,84 | 2,7 | 2,77 | 2,07 |

N1 to N5: biological replicates; NT: Non-Treated; siCTL: Control siRNA; AM: Aberrant Measure.

Melanoma cells proliferation can also be promoted by the level of *TYRP1* mRNA expression.^38^ qPCR analysis show that *TYRP1* mRNA expression level is not significantly affected by the transfection of control, ANXA2 or WDR91 siRNAs (**Supplemental Figure S3C**).

### WDR91 enhancer activity is PS-ASO sequence dependent

We identified WDR91 in melanoma cells using a fully modified TSB ASO and wondered whether similar effects would hold true in a different system (different mechanism of action and cell line). Indeed, it is known that human cancer cell lines have very heterogenous ASO uptake efficiencies. To address this question, we first transfected the U2OS osteosarcoma cell line with siRNAs (control or directed against ANXA2 or WDR91) and evaluated the effect on LNA GMs activity (mRNA knockdown) directed against Metastasis Associated Lung Adenocarcinoma Transcript 1 (*MALAT1*) and Inhibitor of Growth Family Member 2 (*ING2*) RNAs. From these two RNAs, *MALAT1* is a highly studied and conserved nuclear lncRNA and is abundantly expressed in many cells. The experimental workflow is depicted in **Figure 6A**. RT-qPCR in control-siRNA conditions confirmed the efficient downregulation of *MALAT1* and *ING2* mRNAs by their respective GM (**Figure 6B**). No impact was observed regarding either MALAT1 or ING2 ASO-mediated mRNAs knockdowns after ANXA2 downregulation.

**Figure 6.**
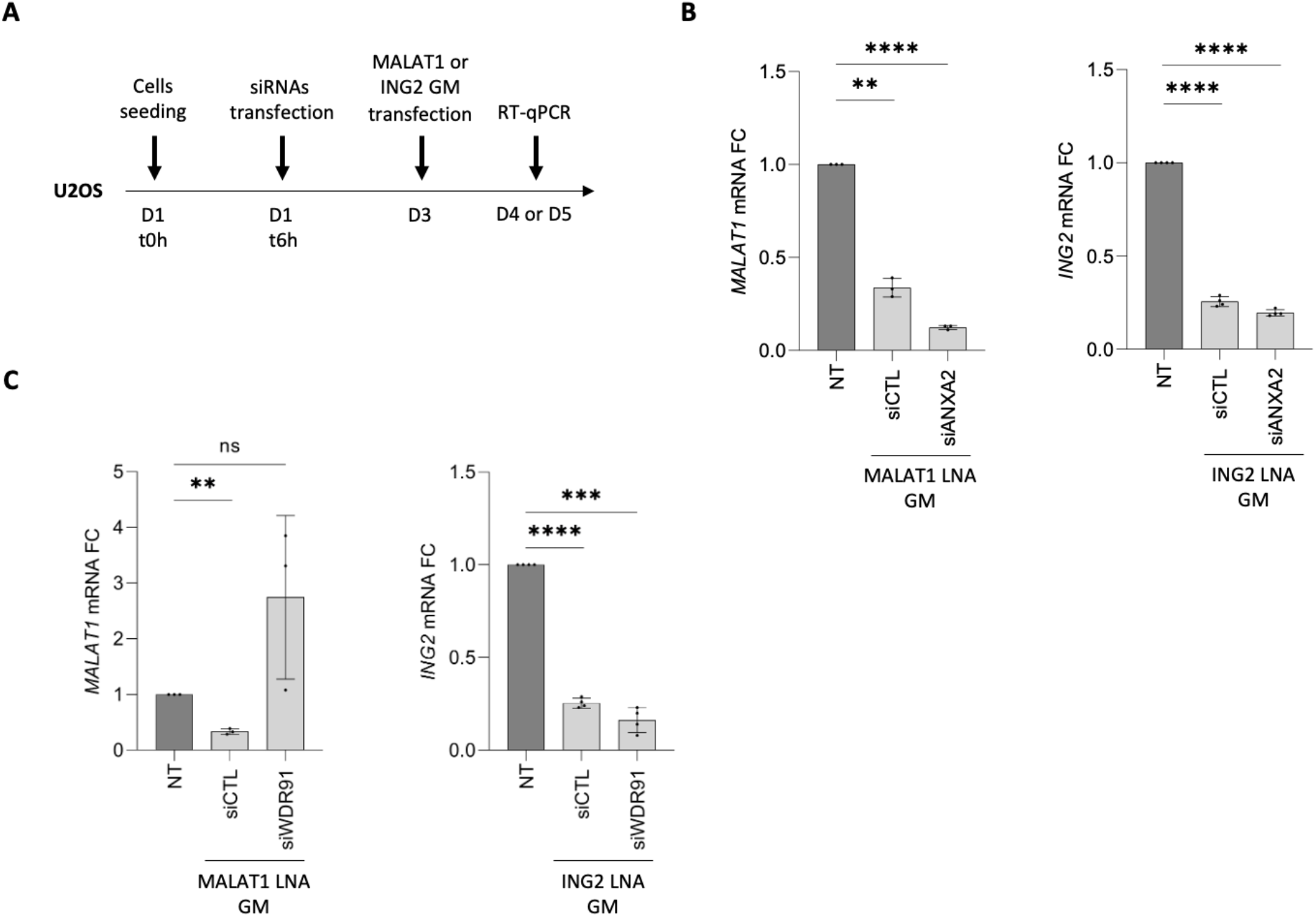
Functional validation of WDR91 in U2OS cells. (A) Workflow of the experimental design (B) *MALAT1* and *ING2* mRNA quantification by RT-qPCR at D4 and D5 respectively, after corresponding GM transfection on non-treated, control or ANXA2 knockdown cells. The relative mRNA fold change was normalized to the non-treated condition (no siRNA transfection). Experiments were performed in independent biological triplicates (MALAT1) and quadruplicates (ING2). Data are presented as mean (SD). Unpaired *t*-test with Welch’s correction were performed. The statistical significance is **P=0,0019 (NT *vs* siCTL); ****P<0,0001 (NT *vs* siCTL; NT *vs* siANXA2) (C) *MALAT1* and *ING2* mRNA quantification by RT-qPCR at D4 and D5 respectively, after corresponding GM transfection on non-treated, control or WDR91 knockdown cells. The relative mRNA fold change was normalized to the non-treated condition (no siRNA transfection). Experiments were performed in independent biological triplicates (MALAT1) and quadruplicates (ING2). Data are presented as mean (SD). Unpaired *t*-test with Welch’s correction were performed. The statistical significance is **P=0,0019 (NT *vs* siCTL); NS= non-significant (P=0,1755); ***P=0,0001 (NT *vs* siWDR91); ****P<0,0001 (NT *vs* siCTL).

We observed that WDR91 knockdown efficiently abrogated MALAT1 ASO activity (**Figure 6C**), confirming the putative role of this protein in ASO potency. However, no such effect was observed on ING2 ASO activity. Both 16-mers MALAT1 and ING2 GMs have the same design and chemistry and only differ by their respective sequence. Considering that proteins interact with ASOs in a sequence dependent manner,^47^ this suggests that WDR91 activity is PS-ASO sequence dependent.

Together, these data highlight for the first time, a set of proteins that can modulate PS-ASO potency in a melanoma cell line, and in a physiological context. Among these candidates, WDR91 is a strong regulator. Its activity has been confirmed in two distinct cell lines but could be ASO-sequence dependent.

### WDR91 does not influence PS-ASOs activity in muscle cells

To evaluate the potency of WDR91 to affect PS-ASOs activity in a different context, we assessed WDR91 ability to influence the activity of a splice-switching ASO used to mediate exon skipping in skeletal muscle cells. ASO-mediated exon-skipping is indeed one of the most promising therapeutic approaches for the treatment of Duchenne muscular dystrophy (DMD). The mechanism of action of ASO in this context is to mask important regulatory splicing signals to exclude the targeted exon and restore the reading frame of the DMD pre-mRNA.^48^ Immortalized human myoblasts from DMD patients are typically used as a model to study dystrophin restoration mediated by splice switching oligonucleotides (SSOs). Here we used immortalized DMD myoblasts carrying a deletion of DMD exon 52 leading to an out-of-frame mRNA and resulting in the absence of dystrophin. The use of an SSO directed against the exon 51 (SSOEx51) allows to restore the open-reading frame of the dystrophin mRNA.

To study the effect of WDR91 on SSOEx51 activity, the immortalized KM571 cells were first transfected with siRNAs to knockdown WDR91 or ANXA2 expression (**Figure 7A**). Seventy-two hours later cells were treated with the SSOEx51 (LNA/2’OMe PS mixmer) to study exon 51 skipping efficiency in a context where WDR91 or ANXA2 expression was reduced. RNA was extracted to first check knockdown efficiency, and we confirmed that siRNAs treatment resulted in a strong knockdown (>70 %) of *WDR91* and *ANXA2* respective mRNAs (**Figure 7B**). Exon skipping levels were then compared and normalized to the siRNA control group (siCTL). We noted a significant reduction in the exon skipping efficacy when ANXA2 expression was reduced (**Figure 7C**). However, this effect was not observed with WDR91. Similar results were obtained with a different SSO chemistry (2MOE PS) (**Supplemental Figure S4**), indicating that WDR91 is not a modulator of PS-ASOs in this system.

**Figure 7.**
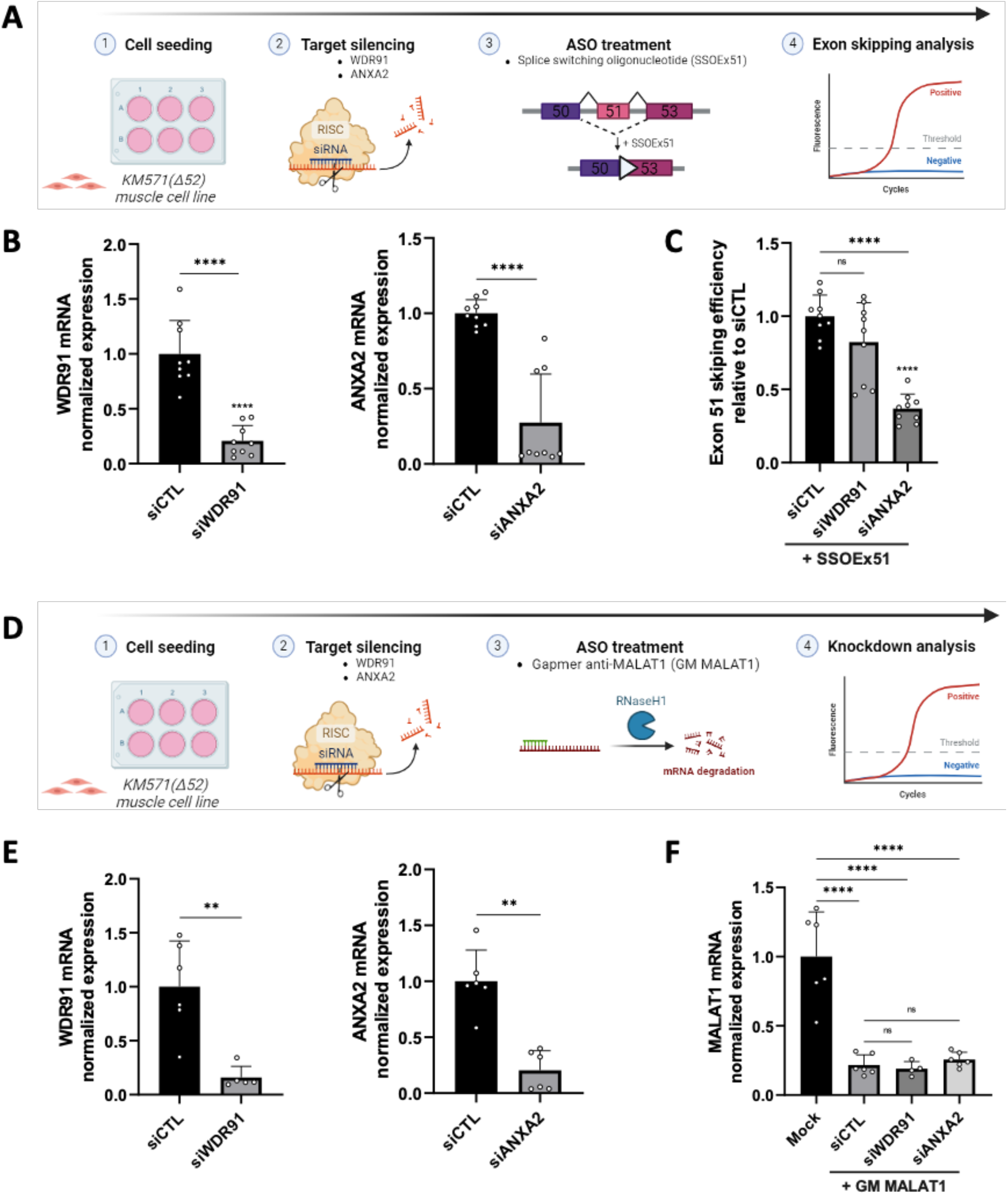
Functional validation of WDR91 in an immortalized DMD muscle cell line (KM571). (A) Experimental setup for the double transfection protocol of siRNAs and SSOs in KM571 (B) *WDR91* and *ANXA2* mRNAs silencing efficiency (siCTL: Control siRNA) (C) *WDR91* and *ANXA2* mRNAs silencing effects on exon 51 skipping efficiency of the SSOEx51 (D) Experimental setup for the double transfection protocol of siRNAs and a GM against MALAT1 (GM MALAT1) in KM571 (E) *WDR91* and *ANXA2* mRNAs silencing efficiency (siCTL: Control siRNA) (F) *WDR91* and *ANXA2* mRNAs silencing effects on MALAT1 knockdown induced by the GM MALAT1. Results are shown as the average of at least four independent biological replicates. Data are presented as mean (SD). The relative mRNA fold change was normalized to the appropriate control condition (Mock or siCTL). Unpaired *t*-test with Welch’s correction were performed. The statistical significance is ** P <0,005; ****P<0,00005.

To determine whether this effect was specific to SSOs or to KM571 cells, we used a different type of ASO to clarify the role of WDR91 in these cells. As described above, we first transfected the KM571 cells with siRNAs targeting *WDR91* or *ANXA2* mRNAs and then with a GM against *MALAT1* mRNA (GM MALAT1) (**Figure 7D**). Here again, siRNA-induced knockdown of *WDR91* and *ANXA2* was very efficient (**Figure 7E**). When quantifying the levels of *MALAT1* mRNA, we observed no significant differences between the siRNA treatment (control, ANXA2 or WDR91) (**Figure 7F**) indicating that WDR91 or ANXA2 knockdown did not affect GM MALAT1 activity. These results suggest that WDR91 likely does not contribute to PS-ASO activity in KM571 cells, regardless of the type of PS-ASOs used.

## DISCUSSION

Despite many clinical successes, the precise mechanisms of ASOs uptake and trafficking remain to be fully understood. In this study, we used a genome-wide CRISPR inhibition screen to identify without *a priori* proteins involved in PS-ASO trafficking, release and productive activity in a 501Mel cell line. Our CRISPR knockout screening strategy has enabled us to discover functionally involved proteins, using a 3^rd^ generation ASO and preserving the integrity of intracellular membranes and organelles. Moreover, the miRNA displacement mediated by a Target Site Blocker ASO represents a relevant assay to investigate the proteins involved in ASOs efficiency and which has been validated *in vivo*.

This study revealed a set of 95 potential activators and 55 inhibitors. Among the activators, our assay highlighted the Rab5C and COG8 proteins, which are known positive regulators of ASOs trafficking and productive activity.^35,36^ ANXA2, which is another known positive regulator, was not found in the list of candidates after application of our filters, which also shows that this type of CRISPR screen can be subject to false negatives.

In this study, we focused on WDR91 protein as a candidate of interest. This protein has a strong biological relevance as it is known to participate in the early-to-late endosome conversion and its knockdown showed the most significant inhibition of tcT3 anti-proliferative activity in our validation experiments.

We confirmed that WDR91 was a positive modulator of PS-ASO activity in two different assays (cell density assay and mRNA knockdown performance), with different cell models (501Mel and U2OS) and ASO chemistries (tcDNA TSB and LNA GM), a landmark result of this project. The impact of WDR91 was not validated in all the systems studied. Indeed, although our results are similar when comparing the ASO activity of WDR91 in 501Mel and U2OS cells, they tend towards an ASO sequence-related activity. In addition, WDR91 did not have any effect with a splice-switching ASO in muscle cells. Thus, as is the case for ANXA2, WDR91 may have an ASO-promoting activity that is also cell line-dependent. Indeed, it is known that identified proteins involved in ASO trafficking and productive activity might not have the same impact depending on the cell line (probably due to differences in high isoform co-expression or the establishment of compensatory proteins or pathways). The same observation was made, for example, in a study evaluating Rab5C, a well-characterized modulator of ASO activity.^49^

Ultimately, the results of our validation experiments on various cell systems and ASO chemistries underline the need for caution when identifying an ASO modulator in a specific context, as it may not be applicable to all systems.

Interestingly, other studies have highlighted WDR91 as an important mediator of the release and efficacy of internalized reoviruses or antibody-drug conjugates in cells during the early endosome maturation or from the late endosomes.^50,32^ Therefore, this study tends to further reinforce the key role of endosomal proteins in productive ASO trafficking and activity.

One of the limitations of our study is that genome-wide CRISPR screening was performed in a single replicate. Rigorous filters were used to identify robust ASO activity modulators. We have chosen to focus on a few selected proteins that promote internalized ASO activity, but it would also be interesting to validate other identified proteins as well as the potential inhibitors to confirm other candidates in 501Mel cells, as well as in other melanoma cell lines with different states of cell differentiation. Furthermore, and given the specificity of this cell type with regard to melanosome biogenesis and transport, involving numerous traffic processes,^51^ it would be of interest to further evaluate WDR91 in other cell lines as it has been done in U2OS and KM571 cells.

Analysis of the gene set enrichment also suggested new avenues to explore. Looking at the inhibitory proteins, some of them were associated with the endosomal cellular compartment. ATP6V0D2, for example, is one of the strongest inhibitor candidates in our selection and is involved in the acidification and pH maintenance in certain intracellular compartments.^52^ Some activator candidates might be related to actin cytoskeleton regulation. From the related proteins, CDC42 and MAP2 are known to be involved in cellular trafficking disorders.^53^ It is therefore relevant to consider that a defect in these proteins could also have an impact on intracellular trafficking and ASO release in melanoma.

Ultimately, our results contribute to a better characterization and identification of protein modulators of ASO activity, providing valuable information for potential new therapeutic avenue. Potent ASO modulators validated *in vitro* could be inhibited or overexpressed in *in vivo* experiments to assess and confirm an increase in ASO potency. Acting at the level of these modulators or directly on ASO-modulators interaction could result in more productive ASO activity, triggering an exacerbated inhibitory effect and an overall increase in the therapeutic index.

## MATERIALS AND METHODS

### Cell lines and lentiviral library

501Mel and U2OS cells were acquired from ATCC. Before experiments, all cells were regularly tested for absence of mycoplasma. Culture and sub-culturing were performed in RPMI-1640 (Gibco) for 501Mel and Mc Coy’s 5A (Gibco) for U2OS, supplemented with 10% fetal bovine serum (FBS) (Gibco), 2mM L-glutamine (Gibco) and 1% penicillin/streptomycin, (Gibco) and cultured at 37°C in a humidified incubator with 5% CO2. Transfections and infections were performed in the same media but without antibiotics and with a decomplemented serum (30 minutes at 56°C). Immortalised myoblasts carrying a deletion of DMD exon 52 (KM571) were obtained from the platform for immortalization of human cells from the Institut de Myologie. KM571 cells were grown in a 50/50 mix of skeletal muscle cell growth medium (Promocell, Germany) and F-10 (Thermo Fisher Scientific, USA) with 10% fetal bovine serum, and 1% penicillin-streptomycin (100 U/mL).

### Antisense oligonucleotides

miRCURY LNA^TM^ Target site blockers TSB-C1 (ACTTTATTACAATCAT) and TSB-T3 (ACACAGTGGCAAACAC) were designed and obtained from Qiagen. tcC1 and tcT3 were synthesized from SQY Therapeutics (Montigny le Bretonneux) as fully modified tcDNA-PS ASOs. The TSB-C1 and tcC1 controls have no significant match to any annotated human 3’UTR. MALAT1 (CGTTAACTAGGCTTTA) and ING2 (TACGTTGGCTTGTTCA) LNA

Gapmers were designed and obtained from Qiagen. For the KM571 experiments, all ASOs were obtained from Eurogentec. Two chemistries of the splice switching oligonucleotide targeting exon 51 (SSOEx51 previously described)^54^ were used: LNA/2’OMe PS mixmer and full 2MOE PS. MALAT1 GM is a 2MOE PS (previously described).^37^ All ASOs were reconstituted in nuclease-free water (Invitrogen).

### siRNAs

siRNAs pools were purchased from Santa Cruz (Negative Control A sc-37007; WDR91 sc- 89398; WBSCR16 sc-89899; FH sc-105377; ELMOD1 sc-77261; TADA2B sc-156169; ANXA2 sc-270151)

### Generation of 501Mel knockout library

Transduction of mycoplasma negative 501Mel cells at 60% confluence was performed in T175 flasks (standard - Sarstedt) with the Brunello knockout pooled lentiviral library from Addgene (# 73179-LV) and in RPMI-1640 decomplemented medium (Gibco) without antibiotics. The Brunello pooled lentiviral library contains 76,441 single guides RNAs (sgRNAs), targeting 19,114 coding genes (4 sgRNAs per gene). A multiplicity of infection (MOI) of 0.4 was selected to ensure that every single cell is infected with only one sgRNA. Twenty-four hours later, the infectious medium was replaced by a fresh complete medium. After 48h, infected cells were positively selected with 2.5µg/mL puromycin (InvivoGen) for 7 days. Following selection and expansion of the surviving cells, around 350 million cells were recovered. A dry pellet of 100 million cells was prepared for gDNA extraction and sequencing. The remaining cells were used for the subsequent reverse transfection.

### RNA interference, ASO treatments and cell density assay

#### 501Mel parental cells

Reverse-transfections of siRNAs were performed in 60mm dishes (standard - Sarstedt) with 224,000 cells per dish and for 48h in RPMI-1640 decomplemented medium without antibiotics. Briefly, 1mL lipoplexes layers were formed with Lipofectamine 2000 (Invitrogen) and the siRNA concentrations were set to 30nM in a final 3mL volume after addition of the cells on top of the lipoplexes. For all the transfections, lipoplexes were formed in OptiMEM^®^. After 16h incubation, the medium was replaced by a fresh complete medium without antibiotics. After 48h, the cells were recovered, counted with a Neubauer counting chamber and a second reverse-transfection was performed in 96-well plates (standard F- Sarstedt) with 3000 cells per well and with either the siRNA alone (at 30nM final concentration) or a combination of siRNA (30nM) and tcT3 or LNA T3 at a final concentration of 50nM in a final volume of 150µL (100µL of cells on top of 50µL of lipoplexes). After 16h incubation, the medium was replaced by a fresh 150µL complete medium without antibiotics. After 4 days following to the second transfection, the cell density was measured with 0.5% crystal violet (10 minutes at RT, followed by three PBS washes, 100µL of 100% EtOH solubilization and absorbance reading at 590nm using a Tecan Infinite F200 Pro reader).

#### U2OS cells

Reverse-transfections of siRNAs were performed in 12-well plates (standard - Sarstedt) with 50,000 cells per well and for 48h in decomplemented Mc Coy’s 5A medium without antibiotics. Cell density was optimized so these cells are efficiently transfected between 20 and 30% confluence. The procedure was the same as described above except that the siRNA transfection was performed with RNAiMax (Invitrogen) and 6h post seeding. After 48h, the medium was replaced by 0.4mL of a fresh decomplemented Mc Coy’s 5A medium without antibiotics and a reverse-transfection with Lipofectamine 2000 was performed in the same plate with the MALAT1 or ING2 Gapmer (at 30nM final concentration) in a final volume of 0.8mL (0.4mL of lipoplexes on top of the cells). After 24h following to the second transfection, total RNA was extracted for further RT-qPCR analysis.

#### 501Mel knockout library

Reverse-transfections of tcT3 and C1 were performed in 150mm dishes (standard - Sarstedt) with 1.5 million knockout cells per dish and for 72h in RPMI-1640 decomplemented medium without antibiotics. Briefly, 10mL lipoplexes layers were formed with Lipofectamine 2000 (Invitrogen) and the respective ASOs concentrations were set to 50nM in a final 24mL volume after addition of the cells on top of the lipoplexes. For the transfections, lipoplexes were formed in OptiMEM^®^ and the mycoplasma-negative knockout cells have been sub-cultured 24h before in T175 flasks (standard - Sarstedt). The cells confluency at the day of transfection was maintained between 50 and 80%. After 7h of transfection, the medium was replaced by a fresh complete medium without antibiotics. To ensure a coverage of 400 cells per guide RNA in our library, a total of 32 million cells was transfected with either the tcT3 or C1. Thus, 22 dishes were used per oligonucleotide transfection. After 72h of transfection, surviving cells were recovered and pellets of 100 million cells were kept at -80°C until further processing.

#### KM571 cells

Cells were plated in 6-well plates (300,000 cells per well) and transfected the same day with 20nM of siRNA with Lipofectamine RNAiMAX according to the manufacturer’s instructions (Thermo Fisher Scientific, USA). Twenty-four hours after the first transfection, proliferation medium was changed for differentiation medium (DMEM, 2% horse serum, 1% penicillin / streptomycin (100 U/mL)) to induce myotubes formation. After 3 days of differentiation, cells were treated with 100nM of the SSOEx51 for 24h with Lipofectamine LTX according to the manufacturer’s instructions (Thermo Fisher Scientific, USA). For the transfection of the GM MALAT1, similar protocol was used, except that KM571 cells were only treated during proliferation and that the GM MALAT1 treatment was performed for 48h.

### NGS sequencing library preparation

Frozen pellets of 100 million transfected surviving cells (and from non-treated knockout library) in 50mL poplypropylene tubes (Sarstedt) were thawed on ice for 20 minutes and lysed at room temperature. Genomic DNA was purified according to the manufacturer’s instructions (NucleoBond HMW DNA, Macherey-Nagel). A total of 280µg of genomic DNAs was used for PCR amplification of sgRNA sequences, using the Herculase II Fusion polymerase (Agilent). Illumina P5- and P7-barcoded adaptors were used, and PCR amplification and quality controls have been carried out as described by Zhang laboratory (Joung *et al*, 2017). Amplified PCR products were purified using AMPure XP beads (Beckman Coulter) according to the manufacturer’s instructions, eluted in 14µL of nuclease-free water (Invitrogen) and stored at - 20°C until Bioanalyzer quality control (TapeStation 4200, Agilent) was used to measure the size, quality and concentration. The pooled samples were sequenced using Illumina NGS sequencing platform (Novaseq6000 – CurieCoreTech ICGex Platform).

### NGS data analysis

MAGeCK (v0.5.9.4) was used to extract sgRNA read counts from the samples fastq.gz files (mageck count program with the --norm-method median option). For each sgRNA, we transformed the read counts onto M (log ratio) and A (mean average): M = log2(sgRNA_count_in_TRT+1) - log2(sgRNA_count_in_CTL+1), and A = mean (count_TRT, count_CTL) in log2 scale. Fold-changes between TRT and CTL conditions were also computed from the M values of each sgRNA.

### RNA extraction and RT-qPCR analysis

Total RNA form cell lysates was obtained using a NucleoSpin^®^ RNA kit (Macherey-Nagel), according to the manufacturer’s instructions and with on-column DNAse treatment. After Nanodrop evaluation of purity and concentration, 500ng of RNA was used for reverse- transcription (MultiScribe RT – Applied Biosystems). Obtained cDNA was diluted to 2.5ng/µL and used as template for qPCR using a QuantStudio5 system (Applied Biosystems). SYBR Green (PowerSYBR^®^ Green PCR Master Mix – Applied Biosystems) was used according to the manufacturer’s instructions. For each sample measurement, 3 technical replicates were performed. TBP was used as internal standard. TYRP1 and ING2 primers were purchased from Eurogentec. % knockdown was calculated as 100*(1-2^-ddCt^). Total RNA was isolated from cultured human myoblast cells using TRIzol reagent, according to the manufacturer’s instructions (Thermo Fisher Scientific, USA). Aliquots of 1 µg of total RNA were used for RT- PCR analysis. The cDNA synthesis was carried out with LunaScript™ RT SuperMix Kit (New England Biolabs). Exon 51 skipping was measured by TaqMan qRT-PCR, using TaqMan assays that were designed against the exon 51-53 or exon 50-53 templates using the Prime Time qPCR probe assays (Integrated DNA Technology) (Assay Ex51-53 - forward : 5’- CAGGTTGTGTCACCAGAGTAA-3’; reverse 5’-TGAGTGGAAGGCGGTAAA-3’; probe : 5’-/56-FAM/TGACCACTA/ZEN/TTGGAGCCTCTCCTAC/3IABkFQ/-3’ and assay Ex50-53: forward: 5’-ACTGATTCTGAATTCTTTCAAAGGC-3’; reverse: 5’- TTCAAGAGCTGAGGGCAAAG-3’; probe: 5’- /56-FAM/ACCTAGCTC/ZEN/CTGGACTGACCACT/3IABkFQ/-3’) . gBlocks Gene Fragments (Integrated DNA Technology) from human exon 49-54 and human exon 49-54 Delta 52 were used as standards for DNA copy number. 70 ng of cDNA was used as input per reaction, and all assays were carried out in triplicate. Assays were performed under fast cycling conditions on a Bio-Rad CFX384 Touch Real-Time PCR Detection System. For a given sample, the copy number of skipped product (exon 50-53 assay) and unskipped product (exon 51-53 assay) was determined using the standards Ex49-54 and Ex49-54 Delta52, respectively. Exon 51 skipping was then expressed as a percentage of total dystrophin. Knockdown analysis was performed by using the appropriate probes for the studied targets and were purchased from Integrated DNA technologies (IDT) (MALAT1 Hs.PT.58.26451167.g; WDR91 Hs.PT.58.39523053 and ANXA2 Hs.PT.56a.40676274). The expression of the different targets was then normalized to a reference gene (GAPDH Hs.PT.39a.22214836).

### Protein isolation and Western blotting

Total protein extracts were obtained by resuspension of PBS-washed cell pellets in 40µL of ice-cold RIPA buffer (Tris-HCl 50 mM, ph7.4, NaCl 150 mM, Sodium Deoxycholate 0.5%, Triton 1%, SDS 0.1%), including protease inhibitor cocktail (Sigma-Aldrich, P8340) and centrifugation at 16,000xg for 15min at 4°C. Proteins quantification from the cleared lysates was performed using a Pierce^TM^ BCA Assay kit (Thermo scientific). After 4-12% SDS-PAGE in NuPAGE MES buffer (Invitrogen), proteins were transferred onto PVDF membranes with a turbo transfer system (BIO-RAD) for 10 minutes at 25V and 2.5A. After transfer, the membranes were washed in Tris-buffered saline and 0.1% Tween 20 (TBST) and blocked for 30 minutes at room temperature in a solution of TBST + 5% w/v non-fat dry milk. Blocked membranes were washed three times in TBST for 5 minutes and diluted primary antibody was incubated at 4°C overnight with shaking. The diluted HRP-conjugated secondary antibody was added for 1h at room temperature with shaking, after three TBST washes of 5 minutes. Imaging was performed after ECL reagent addition and chemiluminescence detection with G:Box Chemi XX6 imager from Syngene. ANXA2 antibody was purchased from Santa Cruz (sc- 28385).

### Statistical analysis

GraphPad Prism software (San Diego, CA, USA) version 9.5.1 was used for statistical analysis and graphs design. Each data is presented as mean of at least three independent biological replicates with standard deviation (SD). P-values were calculated by two-sided unpaired *t*-tests with Welch’s correction and differences were statistically significant at *p ≤ 0.05; **p ≤ 0.01; ***p ≤ 0.001; ****p ≤ 0.0001. For exon skipping experiments, normal distribution of samples was assessed by using Shapiro-Wilk test. Comparisons of statistical significance were assessed by one-way ANOVA parametric test or Kruskal-Wallis non-parametric test and followed by appropriate post-hoc tests if applicable.

## DATA AVAILABILITY STATEMENT

Data are available upon request to the corresponding author.

## ACKNOWLEDGMENTS

The authors thank the SFR Biosit L3 platform of Rennes University for providing the required facility for performing the lentiviral infections. The authors also thank the Institut Curie (Genetic screen CRISPR’it and ICGex Core Facilities) for performing the NGS sequencing, SQY Therapeutics for providing the tcDNA-ASO molecules and the platform for the immortalization of human cells from the Institut de Myologie for the human-derived cells. This study received financial support from the Agence Nationale de la Recherche (ANR).

## AUTHOR CONTRIBUTIONS

G.M., R.P., D.G., A.Go. and A.Ga. conceived and analyzed the experiments. G.M. and A.Ga. performed the experiments. M.A. analyzed the NGS raw data. A.M. and A.B. helped in the library preparation. G.M. wrote the manuscript. All the authors reviewed the manuscript.

---

**Supplemental Figure S1**. **Gene set enrichment analysis on activator candidates.** Gene set enrichment analysis using PANGEA (Pathway, Network and Gene-set Enrichment Analysis) allowed to classify the activator candidates based on the cellular component gene ontology. Each gene set is classified according to its highest value (log2 fold change). In green are framed the proteins that are part of the peroxisome cellular component. In red are framed the proteins that are part of the mitochondrion cellular component. In purple are framed the proteins that are part of the actin cytoskeleton cellular component. The color code and corresponding lines highlight common candidates between each corresponding cellular component.

**Supplemental Figure S2**. **Gene set enrichment analysis on inhibitor candidates.** Gene set enrichment analysis using PANGEA (Pathway, Network and Gene-set Enrichment Analysis) allowed to classify the inhibitor candidates based on the cellular component gene ontology. Each gene set is classified according to its highest value (log2 fold change). In green are framed the proteins that are part of the Golgi apparatus cellular component. In red are framed the proteins that are part of the microtubule cytoskeleton cellular component. In purple are framed the proteins that are part of the endosomal cellular component. The color code and corresponding lines highlight common candidates between each corresponding cellular component.

**Supplemental Figure S3.** (A) Western blot analysis of ANXA2 protein expression at 48h and 72h on non-treated 501Mel cells and after ANXA2 or control siRNA reverse transfections. NT: non-treated; siANXA2: ANXA2 siRNA; siCTL: Negative control siRNA. Alpha-tubulin was used as a loading control (B) Evaluation and quantification of the candidate’s knockdown efficiency by RT-qPCR on total RNAs coming from non-treated (NT) or siRNA transfected 501Mel cells. Data are calculated from three technical replicates. % knockdown (KD) was calculated as 100*(1-2^-ddCt^). TBP was used as housekeeping gene (C) TYRP1 mRNA quantification by RT-qPCR on total RNAs coming from non-treated (NT) or siRNA transfected 501Mel cells (CTL: control siRNA). Results are shown as the mean of at least three independent biological replicates (SD).

**Supplemental Figure S4**. **WDR91 do not affect the activity of a methoxyethyl chemistry splice switching oligonucleotide in DMD muscle cells.** SSOEx51 used in these experiments is an 2MOE PS. (A) *WDR91* and *ANXA2* mRNAs silencing efficiency (siCTL: Control siRNA) (B) *WDR91* and *ANXA2* mRNAs silencing effects on exon 51 skipping efficacy of the SSOEx51 2MOE PS. Results are shown as the average of at least four independent biological replicates. Data are presented as mean (SD). The relative mRNA fold change was normalized to the appropriate control condition (Mock or siCTL). P-value was calculated by two-sided unpaired t-tests against the Mock transfected or control siRNA condition. The statistical significance is ** P <0.005; ****P<0.00005.

