## Supplementary material for "A Genome-wide CRISPR screen unveils the endosomal maturation protein WDR91 as a promoter of productive ASO activity in melanoma"

### Supplemental Figure S1

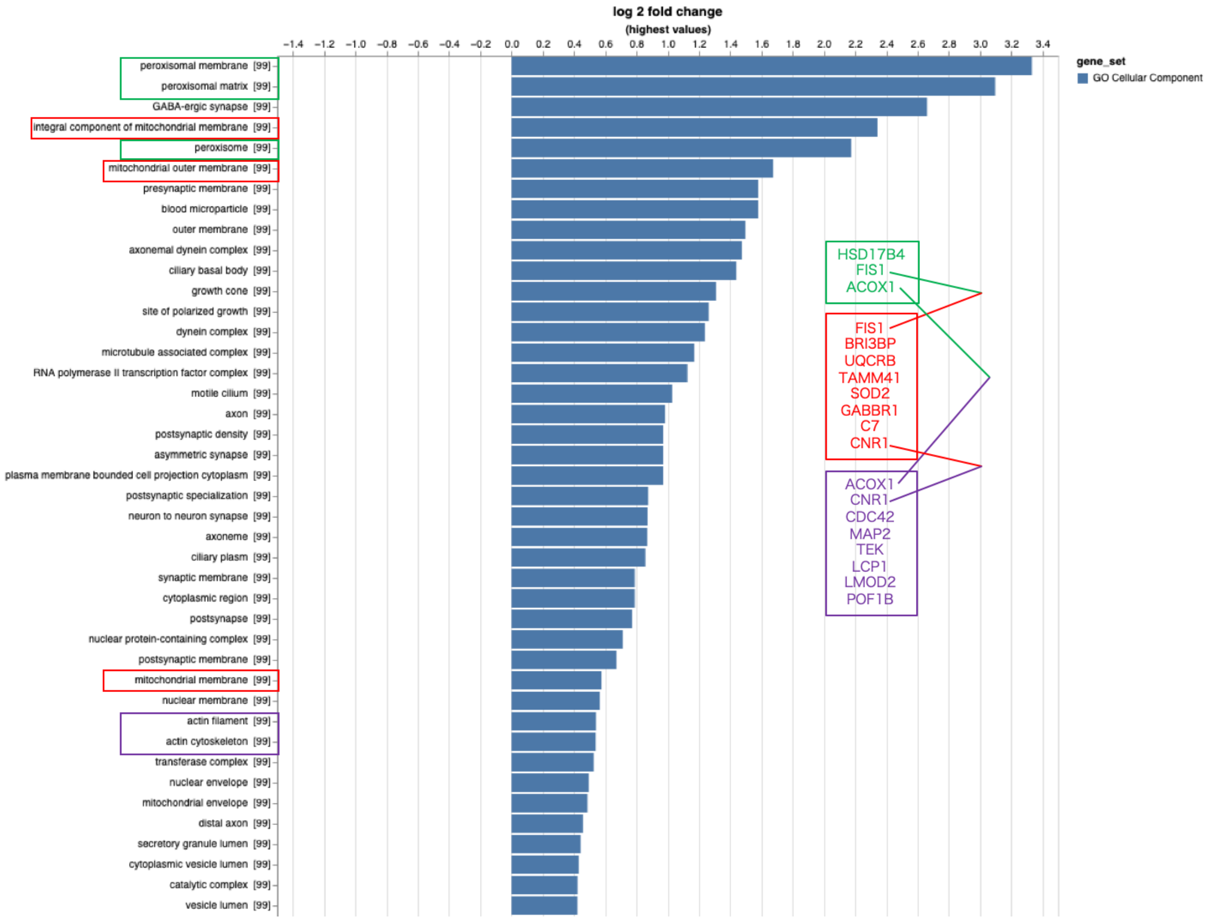

### Supplemental Figure S2

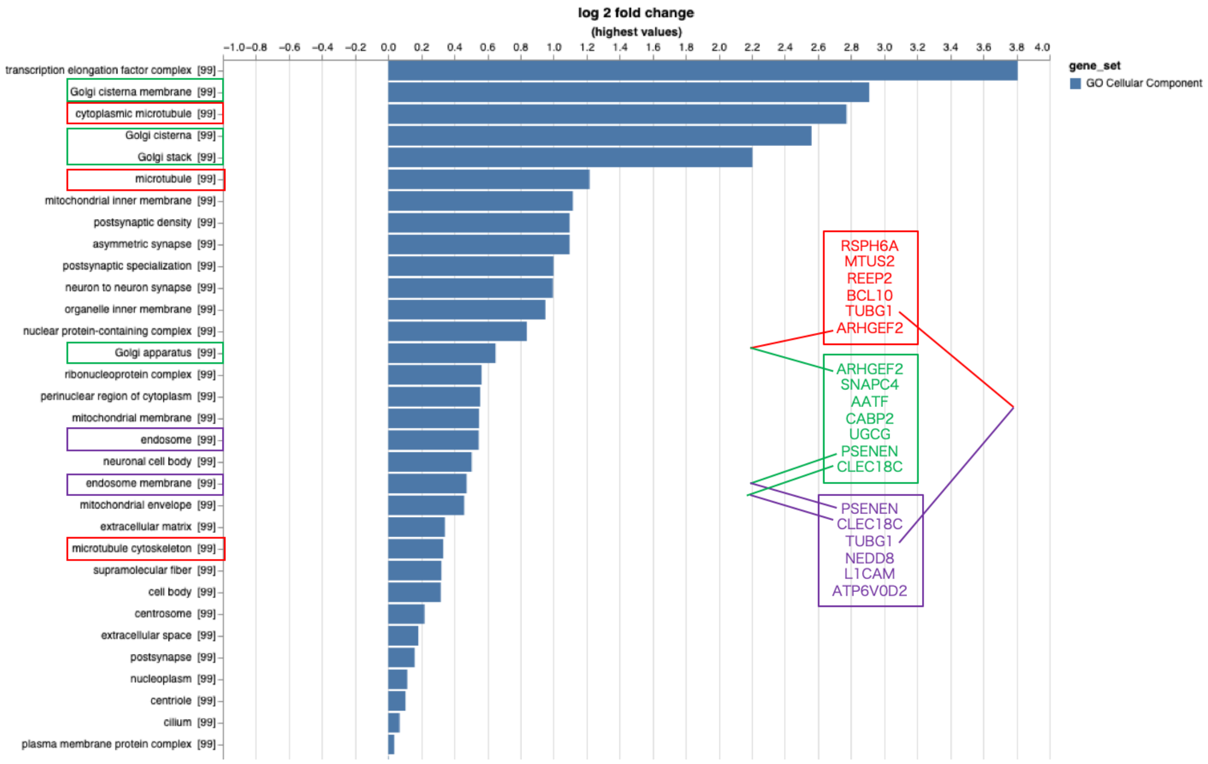

### Supplemental Figure S3

**A**

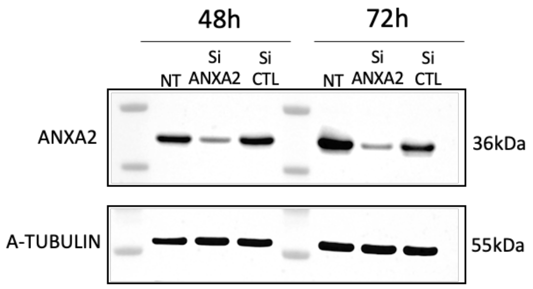


**B**

| siRNA | % KD |
|---|---|
| WDR91 | 50,4 |
| WBSCR16 | 80,4 |
| FH | 93,1 |
| ELMOD1 | 26,6 |
| TADA2B | 84,6 |

**C**

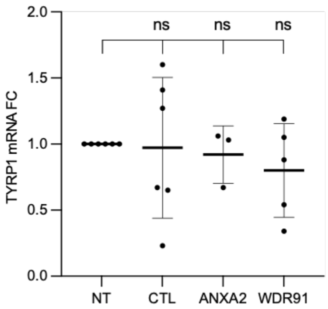

**Supplemental Figure S4**

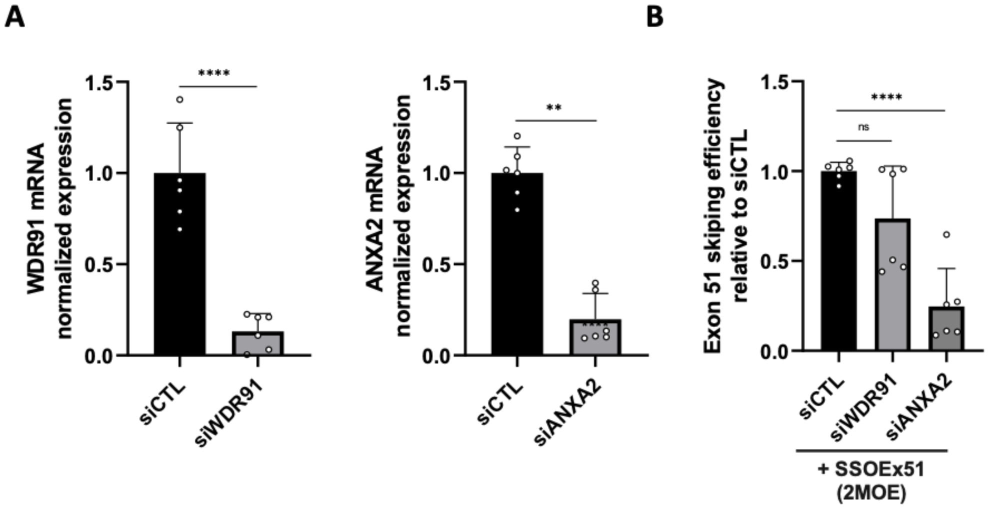
